# Flagellar system translocates T3SS effectors critical for *Salmonella* infection

**DOI:** 10.64898/2026.06.04.729515

**Authors:** Alona Dreus, Paulina Matheova, Martina Spejzlova, Grzegorz Grabe, Jana Schmidtova, Milada Pospíšilová, Tomas Brabec, Ondrej Cerny

**Author notes:** Corresponding author: Ondrej Cerny, Laboratory of Bacterial Virulence, Institute of Microbiology of the Czech Academy of Sciences, Prague, Czech Republic, Videnska 1083, Prague, 142 20, Czech Republic.

## Abstract

Type 3 secretion system (T3SS) injectisomes, the essential virulence factors of many bacterial pathogens, are thought to have evolved from the flagellar T3SS. Despite their high degree of structural similarity, whether flagella can translocate effector proteins typically associated with T3SS injectisomes remains poorly understood. Here, we demonstrate that the flagellar T3SS of *Salmonella* translocates T3SS effectors directly into the host cell cytosol and that this process is critical for productive infection. We show that upon host cell adhesion, flagella were required for efficient *Salmonella* invasion. Fully assembled flagella enabled effector translocation in a process dependent on the flagellar motor. Notably, flagella continued to translocate effectors within the *Salmonella*-containing vacuole until expression of the *Salmonella* pathogenicity island 2–encoded T3SS injectisome was induced. Together, these findings reveal a previously overlooked function of bacterial flagella in host–pathogen interactions and redefine their contribution to *Salmonella* virulence.

## Introduction

Many virulence factors have evolved as tools that enable pathogenic bacteria to invade their hosts, survive within host environments, and disseminate to new tissues. Overlapping functions among individual virulence factors result in redundancy, which enhances the robustness of pathogenicity. As a result, investigation of a given virulence factor often requires deletion of multiple additional factors to reveal its specific contribution to infection^1^. Effector proteins of the type 3 secretion system (T3SS) injectisomes provide a good example of such redundancy^2^.

T3SS injectisomes are essential virulence factors of many extracellular and intracellular bacterial pathogens. These macromolecular machines manipulate biological processes of host cells through the translocation of effector proteins. Structurally, the T3SS injectisome resembles a molecular syringe. The injectisome consists of a cytosolic sorting platform, an inner membrane ring, a rod, an outer membrane ring, an extracellular needle, and a translocon complex inserted into the host cell plasma membrane^3^. Upon the host cell contact, the needle tip mediates the formation of a translocon pore in the host membrane, thus establishing a channel for the translocation of unfolded effector proteins across the inner and outer bacterial membrane and plasma membrane of the target host cell. Translocation of virulence factors is a metabolically costly process, which requires functioning of ATPase machinery^4^. The facultatively intracellular enteropathogen *Salmonella enterica* serovar Typhimurium is often used as a model organism to study the structure and function of T3SS injectisomes.

Virulence of *S.* Typhimurium depends on two distinct T3SS injectisomes encoded on *Salmonella* pathogenicity islands (SPI) 1 and 2. The SPI-1–encoded injectisome is expressed in the gut lumen and is essential for gastrointestinal disease^5^ and for active invasion into host cells^6,7^. Following entry into host cells, *Salmonella* resides within the so-called *Salmonella*-containing vacuole (SCV), where the expression of the SPI-1 injectisome is rapidly downregulated. Following initial establishment of the SCV and its gradual acidification, expression of the SPI-2–encoded injectisome is induced^8,9^. The SPI-2 injectisome is required for bacterial replication within the SCV^10,11^ and contributes to both intestinal^12^ and systemic infections^13^.

To date, approximately 40 effector proteins translocated by the SPI-1 and SPI-2 injectisomes have been identified^14^. While some of these effectors can be translocated by both injectisomes, most are reportedly translocated exclusively by either SPI-1 or SPI-2 apparatuses. Although some of the effectors exhibit overlapping functions, others exert opposing activities in host cells, such as pro- and anti-inflammatory effects. Notably, effectors translocated by the SPI-1 injectisome are delivered in temporally distinct phases of infection to circumvent functional interference between different effector subsets^15^. This suggests that switching from SPI-1– to SPI-2–dependent effector translocation represents a critical regulatory mechanism during infection. However, several effectors such as PipB2 can be translocated by both injectisomes and are involved in SCV maturation; therefore, any interruption in their delivery would pose a significant challenge to the intracellular lifestyle of *Salmonella*^16,17^.

Although the SPI-1 and SPI-2 injectisomes belong to distinct structural families (i.e. the SPI-1 injectisome to the Inv-Mxi-Spa family and the SPI-2 injectisome to the Ssa-Esc family), they share an evolutionary origin and structural similarity with the flagellar T3SS^18^. The core conserved part of all three T3SS is placed in the transmembrane and cytosolic compartments and forms the basal body of the apparatuses. It consists of export apparatus, the surrounding inner membrane (IM) ring and a cytosolic ATPase connected via a stalk to the C-ring^19^. The major structural divergence between the injectisomes and the flagellar T3SS lies in the extracellular appendages: the flagellum contains the rod–hook–filament complex, whereas the virulence-associated SPI-1 and SPI-2 T3SSs possess needle-like structures (Extended Data Fig. 1, Extended Data Table 1)^20^.

While flagellum-dependent motility and chemotaxis are widely recognized as important contributors to bacterial virulence^21,22^, the potential involvement of the flagellar T3SS in effector translocation remains poorly explored. Moreover, little is known about how different virulence-related functions of one structure interact and are co-ordinately regulated during infection.

To test the hypothesis that the *Salmonella* flagellum acts as an additional system for effector delivery during infection, we tracked effector proteins at the level of host-cell translocation and correlate their secretion with other flagellum-related functions. Here we show effector proteins historically associated with *Salmonella* injectisomes are also translocated by the flagellar T3SS, both by the bacteria adhered to host cells from the extracellular space and by the intracellular bacteria residing in SCV. A complete flagellar structure is needed for efficient effector translocation. We also demonstrate that flagella-mediated effector translocation is involved in *Salmonella* invasion into host cells and in the first phase of *Salmonella* intracellular life. Then flagella disassemble in the SCV approximately when the SPI-2 injectisome is induced. The effector translocation by the flagellar T3SS reported here highlights a previously overlooked function of the flagellum in bacterial virulence.

## Results

### *Salmonella* invasion into epithelial cells depends on flagella

Previous studies showed involvement of bacterial flagella in adhesion to and invasion into host cells^23–25^. However, the distinction between adhesion and invasion, and the direct function of flagella in invasion has not been described in detail. To distinguish between extracellular (adherent) and intracellular (invading) bacteria, we infected HeLa cells with *Salmonella* stably expressing GFP, and labelled extracellular bacteria with *Salmonella-*specific antibody thus allowing their discrimination using flow cytometry (Fig. 1A).

**Figure 1.**
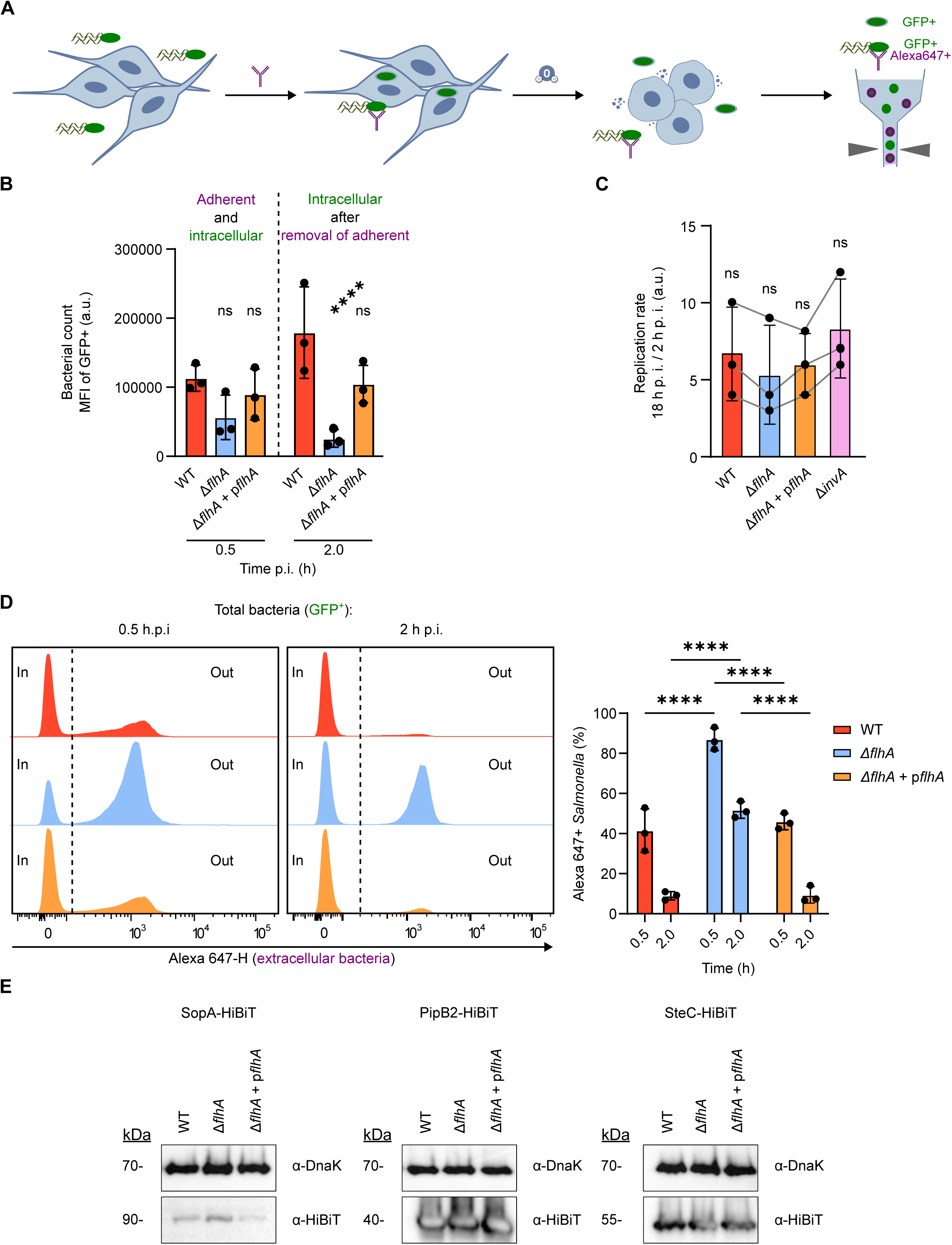
Flagellar effector translocation supports Salmonella invasion into epithelial cells. **A.** Schematic representation of *Salmonella* adhesion and invasion comparison. *Salmonella glmS::*P*trc*-GFP grown to late exponential phase was used to infect HeLa cells for 30 minutes. After removal of nonadherent extracellular bacteria, adherent extracellular *Salmonella* were labelled with primary anti-*Salmonella* antibody and AlexaFlour 647-conjugated secondary antibody. After osmotic lysis of HeLa cells, the extracellular adherent AlexaFlour 647⁺GFP⁺ and intracellular AlexaFlour 647⁻GFP⁺ *Salmonella* was quantified using flow cytometry. **B.** Total amount of *Salmonella* WT *glmS::*P*trc*-GFP, *Salmonella glmS::*P*trc*-GFP Δ*flhA*, and *Salmonella glmS::*P*trc*-GFP Δ*flhA* + p*flhA* (GFP+ irrespective of AlexaFlour 647 signal) was quantified from lysate of infected HeLa cells at 0.5 and 2 h p.i. Data represent mean values from three independent experiments in technical duplicates. ns, p > 0.05; *** p < 0.001 (Two-way ANOVA with Tukey HSD test). **C.** Replication rate of intracellular *Salmonella* WT, Δ*flhA*, Δ*flhA* + p*flhA*, and Δ*invA* (SPI-1 null control), determined by CFU enumeration at 2 and 18 h p.i. Data represent mean values from three independent experiments in technical duplicates; the values from each experiment are connected by grey lines. ns, p > 0.05 (One-sample Wilcoxon test). **D.** Quantification of adherent (Alexa647⁺GFP⁺) and intracellular (Alexa647⁻GFP⁺) bacteria in HeLa cells infected with *Salmonella* WT *glmS::*P*trc*-GFP, *Salmonella glmS::*P*trc*-GFP *flhA::Cm*, and *Salmonella glmS::*P*trc*-GFP *flhA::Cm* + p*flhA* at 0.5 and 2 h p.i. Representative data (left) and mean values from three independent experiments performed in technical duplicates (right) are shown. **** p < 0.0001 (Two-way ANOVA with Tukey HSD test). **E.** Quantification of SopA, PipB2, and SteC protein levels in *Salmonella* using immunoblotting detection. Effector proteins were detected in lysate of *Salmonella* WT, *Salmonella* Δ*flhA*, and *Salmonella* Δ*flhA* + p*flhA* cultures using anti-HiBiT tag antibody. DnaK *Salmonella* protein was used as a loading control. Representative of 3 independent experiments.

We observed no significant difference in HeLa cell-associated bacterial count of *Salmonella* (i.e. adherent to the surface and invaded into HeLa cells) at 0.5 h post infection (p.i.) between *Salmonella* WT, Δ*flhA*, and Δ*flhA* + p*flhA* strains, suggesting that flagella did not function as a major adhesin in this experimental setup (Fig. 1B). In contrast, we detected significantly less *Salmonella* Δ*flhA* at 2 h p.i. when the infected cells were washed with PBS, and the extracellular bacteria were killed with gentamicin (Fig. 1B). We observed no difference in replication rate between intracellular *Salmonella* WT, Δ*flhA*, and Δ*flhA* + p*flhA* strains (Fig. 1C), suggesting that the observed difference in bacterial count resulted from involvement of flagella in *Salmonella* invasion and was not caused by a replication defect.

At 0.5 h p.i., approximately 90% of the HeLa cell-associated *Salmonella* Δ*flhA* mutant population remained extracellular, compared with only 40% of *Salmonella* WT and complemented *Salmonella* Δ*flhA* + p*flhA* bacterial populations (Fig. 1D). At 2 h p.i., the proportion of extracellular *Salmonella* Δ*flhA* bacteria decreased to ∼50%, whereas *Salmonella* WT and Δ*flhA* + p*flhA* complemented strains showed only ∼10% of extracellular bacteria (Fig. 1D). Since the extracellular bacteria were killed by gentamicin before finishing the invasion into the host cells, these results further highlight the importance of flagella for *Salmonella* invasion.

Previous studies have shown that gene deletions can have unintended effects on overall bacterial physiology, such as polar effects or epistasis^26,27^. To test whether deletion of *flhA* affects *Salmonella* virulence in an indirect manner, we examined production of selected effector proteins. Bacteria were cultured under SPI-1– or SPI-2–inducing conditions to assess the expression of the SPI-1 injectisome effector SopA, or PipB2 known to be translocated by both SPI-1 and SPI-2 injectisomes, and SteC that is translocated by the SPI-2 injectisome, respectively. Western blot analysis revealed no differences in effector protein production among *Salmonella* WT, Δ*flhA*, and Δ*flhA* + p*flhA* complemented strains (Fig. 1E). This shows that the *Salmonella* Δ*flhA* invasion defect is not caused by changes in effector expression.

### *Salmonella* T3SS effectors are translocated with hierarchy exceeding SPI-1 and SPI-2 injectisomes

Since flagellar T3SS shows high degree of similarity to both injectisomes^20,28^ (Extended Data Table 1), we hypothesized that flagellar T3SS supported *Salmonella* invasion by delivering injectisome effectors into host cells. To monitor effector translocation, we used the split luciferase NanoLuc described previously^29^. This approach enables sensitive detection of HiBiT-tagged effectors after their translocation into the host cell cytosol, where the HiBiT tag interacts with ectopically expressed LgBiT part of the luciferase enzyme, resulting in a strong luciferase signal (Fig. 2A).

**Figure 2.**
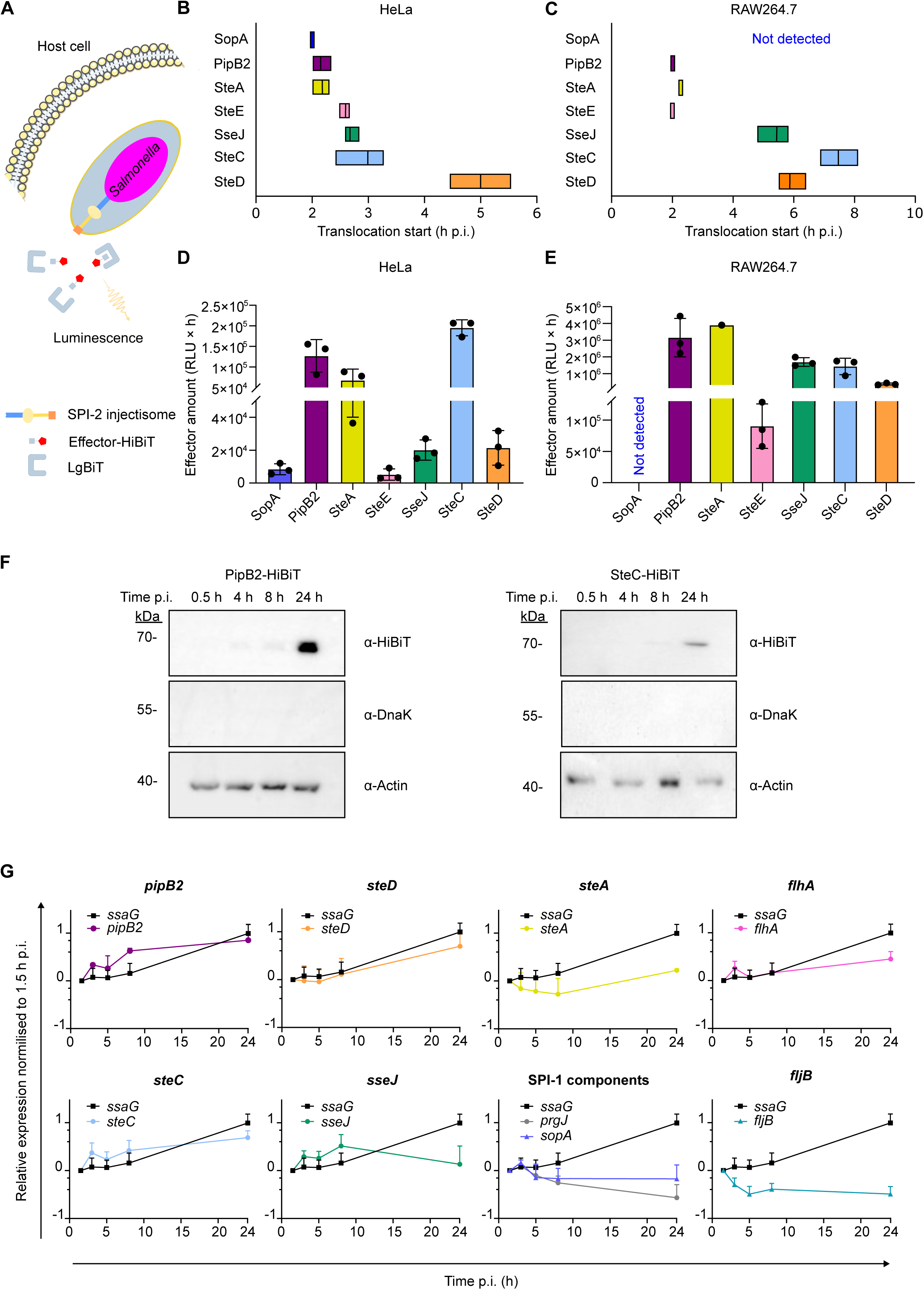
Hierarchy of effector protein expression and translocation. **A.** Schematic representation of LgBiT-HiBiT splitting assay. Following infection of host cells with *Salmonella* strain expressing a HiBiT-tagged effector, extracellular bacteria were removed with extensive PBS washing and remaining extracellular bacteria were killed using gentamicin treatment. Translocation of the HiBiT-tagged effector was monitored as luminescent signal originated from complemented Nano-Luc (*Salmonella*-delivered HiBiT complementing host cell-expressed cytosolic LgBiT). Resulting luminescence recording was started at 2 h p.i. and continued for up to 20 h. **B and C.** Start time of effector translocation into HeLa-LgBiT (B) and RAW264.7-LgBiT (C) cells. Data from three independent experiments performed in duplicates are shown as floating bars with line at mean. Translocation start time was determined by fitting an asymmetric Gompertz model to the luminescence raw data using the script in Appendix A. **D and E.** Amounts of effectors translocated by intracellular bacteria from experiments shown in B and C. Normalised and raw data are shown in Extended Data Fig. 2B-E. **F.** Quantification of PipB2 and SteC translocation using immunoblotting detection. Translocated effector proteins were detected in lysate of HeLa cells infected as in A at indicated time p.i. using anti-HiBiT tag antibody. Anti-DnaK *Salmonella* antibody was used to exclude samples containing lysed bacteria, anti-Actin antibody was used as a loading control. Representative of 3 independent experiments. **G.** Expression profiles of effector proteins in HeLa cells infected with *Salmonella* WT for 1.5, 3, 5, 8, and 24. Data were normalised to the mRNA levels at 1.5 h p.i. for each effector. Expression of *ssaG* was used to allow comparison between individual genes. Values represent the mean of at least three independent experiments performed in duplicate.

To evaluate the limitations of this assay, we first monitored translocation of *Salmonella* HiBiT-tagged effectors into epithelial cells and macrophages constitutively expressing LgBiT. We focused on effectors with well described promoter regions allowing reliable cloning strategy. As expected, translocation of SopA-HiBiT, a SPI-1 injectisome effector, into HeLa-LgBiT cells resulted in an immediate strong luminescent signal that began to decrease soon after measurement initiation, when the extracellular bacteria were removed (Extended Data Fig. 2A). This observation is consistent with the described translocation of SopA by the SPI-1 injectisome^15^. Translocation of the SPI-2 injectisome effector SteD-HiBiT was not detected until after 5 h p.i., as expected from reported time of SPI-2 induction^9^, and the amount of translocated SteD-HiBiT grew steadily to at least 10 h p.i. in HeLa-LgBiT (Fig. 2B, D, and Extended Data Fig. 2B, C). Translocation of several effectors started early after measurement initiation but continued for several hours. Among these, the HiBiT-tagged SteA, SteE, and PipB2, which can be translocated by both SPI-1 as well as SPI-2 injectisomes^30^, were identified (Fig. 2B, D and Extended Data Fig. 2B, C), suggesting that the split NanoLuc assay can be used effectively for the continuous detection of effector translocation by different secretion systems. Interestingly, translocation of SseJ-HiBiT and SteC-HiBiT was detected as early as 3 h p.i., suggesting that these effectors might also be translocated by a secretion system other than the SPI-2 injectisome (Fig. 2B, D, and Extended Data Fig. 2B, C) as described for other effectors previously^30^.

To examine the influence of host cell type on effector translocation, we constructed the RAW264.7-LgBiT macrophage-like cell line. However, we did not detect measurable effector translocation in RAW264.7-LgBiT cells using the NanoGlo substrate used in HeLa-LgBiT and had to use the Endurazine substrate. The Endurazine substrate has longer cell permeability time and is more suitable for prolonged measurements, which makes it optimal for detection of SPI-2 injectisome effector translocation. However, this substrate does not seem to permeate into HeLa-LgBiT cells, thus, different NanoLuc substrates were used for the two cell lines.

The normalised translocation of all effectors followed distinct pattern in RAW264.7-LgBiT (Fig. 2C and Extended Data Fig. 2D, E) from that in HeLa-LgBiT (Fig. 2B and Extended Data Fig. 2B and C). We did not observe the SopA translocation in RAW264.7-LgBiT cells (Extended Data Fig. 2A). However, translocation of PipB2 and SteE was detected from the beginning of measurement suggesting that the Endurazine substrate is suitable for early detection of effector translocation and does not influence the timing of detection. Interestingly, translocation of SseJ and SteC was delayed in RAW264.7-LgBiT in comparison to HeLa-LgBiT cells, whereas SteD was translocated in both cell types at the same time post infection (Fig. 2B to E). Importantly, comparison of effector translocation start times between HeLa-LgBiT and RAW264.7-LgBiT suggest that the hierarchy in effector translocation extends beyond the simple distinction between SPI-1 and SPI-2 injectisome-dependent translocation. Instead, it reveals a more refined hierarchy among effectors that can be translocated by the SPI-2 injectisome.

To further validate effector translocation into host cells and link it with its accumulation, we detected the effectors translocated into the cytosol of HeLa-LgBiT cells using immunoblotting. To reach the amount of signal detectable early after infection, we focused on PipB2-HiBiT and SteC-HiBiT that showed the strongest signal overall (Fig. 2D and E). We observed detectable translocation of PipB2-HiBiT into HeLa cells already at 4 h p.i., while SteC-HiBiT translocation was observable earliest at 8 h p.i. (Fig. 2F), consistent with PipB2-HiBiT detection preceding SteC-HiBiT in the cell luciferase assay. Both effectors accumulated in the host cell cytosol throughout the rest of the experiment (Fig. 2F), suggesting that the luciferase assay reliably detects the onset of effector translocation but not its accumulation in the late time points.

### Expression of effectors within host cells correlates with translocation

While SPI-1 injectisome effectors must be expressed and stored within *Salmonella* prior contact with a host cell to allow immediate response^15^, effector expression by intracellular bacteria responds to conditions within SCV and may be reflected by immediate translocation. To test the sensitivity of HiBiT – LgBiT detection, we correlated translocation signal with effector expression. As expected for the SPI-1 injectisome, qPCR gene expression analysis of *prgJ*, encoding the inner rod protein, and of *sopA* effector gene by WT *Salmonella* showed that it was already low in comparison to the *ssaG* encoding the SPI-2 needle protein at 1.5 h p.i. and further decreased during the whole experiment duration (Extended Data Fig. 2F). In contrast, expression of the exclusively SPI-2 injectisome effector SteD was low at the beginning of the experiment and started to grow only after 5 h p.i. (Fig. 2G and Extended Data Fig. 2F), consistent with the effector translocation (Fig. 2B and C). Effectors that started their translocation early and were translocated for a prolonged period were expressed in a pattern comparable to each other and in similar amounts throughout the experiment (Fig. 2G). Importantly, expression of *ssaG*, encoding the SPI-2 injectisome needle protein, started to grow strongly after 5 h p.i. and increased throughout the experiment, consistently with the pattern of *steD* expression (Fig. 2G). This suggests that between the end of SPI-1 injectisome-dependent translocation and the beginning of SPI-2 injectisome-dependent translocation, the effectors could be translocated by an alternative route such as the flagellar T3SS.

### Flagellar T3SS is involved in translocation of injectisome effectors

To examine the involvement of flagellar T3SS in effector delivery, we measured effector translocation by *Salmonella* mutant strains lacking the inner membrane export gate protein SctV of each T3SS encoded by *flhA* (for flagella), *invA* (in SPI-1), or *ssaV* (in SPI-2) (Fig. 3A). In macrophage-like RAW264.7 cells, where the SPI-1 injectisome is not needed for *Salmonella* internalisation, Δ*invA* mutant lacking SPI-1 activity displayed a pronounced delay in translocation of both PipB2-HiBiT and SteC-HiBiT (Fig. 3B, C and Extended Data Fig. 3A to D), suggesting that the SPI-1 injectisome is crucial for initial phase of PipB2 and SteC translocation. Interestingly, the Δ*invA* mutant showed a significant decrease in the amount of translocated PipB2-HiBiT (Fig. 3D), while the overall amounts of translocated SteC-HiBiT, that is translocated later (Fig. 2C), was not influenced (Fig. 3E), suggesting translocation of SteC might be compensated by the SPI-2 injectisome later during the infection. The SPI-2 translocation-deficient Δ*ssaV* mutant showed significantly decreased SteC-HiBiT translocation (Fig. 3E) that was accompanied by only a small early peak of effector translocation (Extended Data Fig. 3D), while PipB2-HiBiT translocation was not significantly influenced by Δ*ssaV* mutation during RAW264.7-LgBiT infection (Fig. 3D). The Δ*flhA* mutation did not significantly influence effector translocation into RAW264.7 macrophages (Fig. 3B to E).

**Figure 3.**
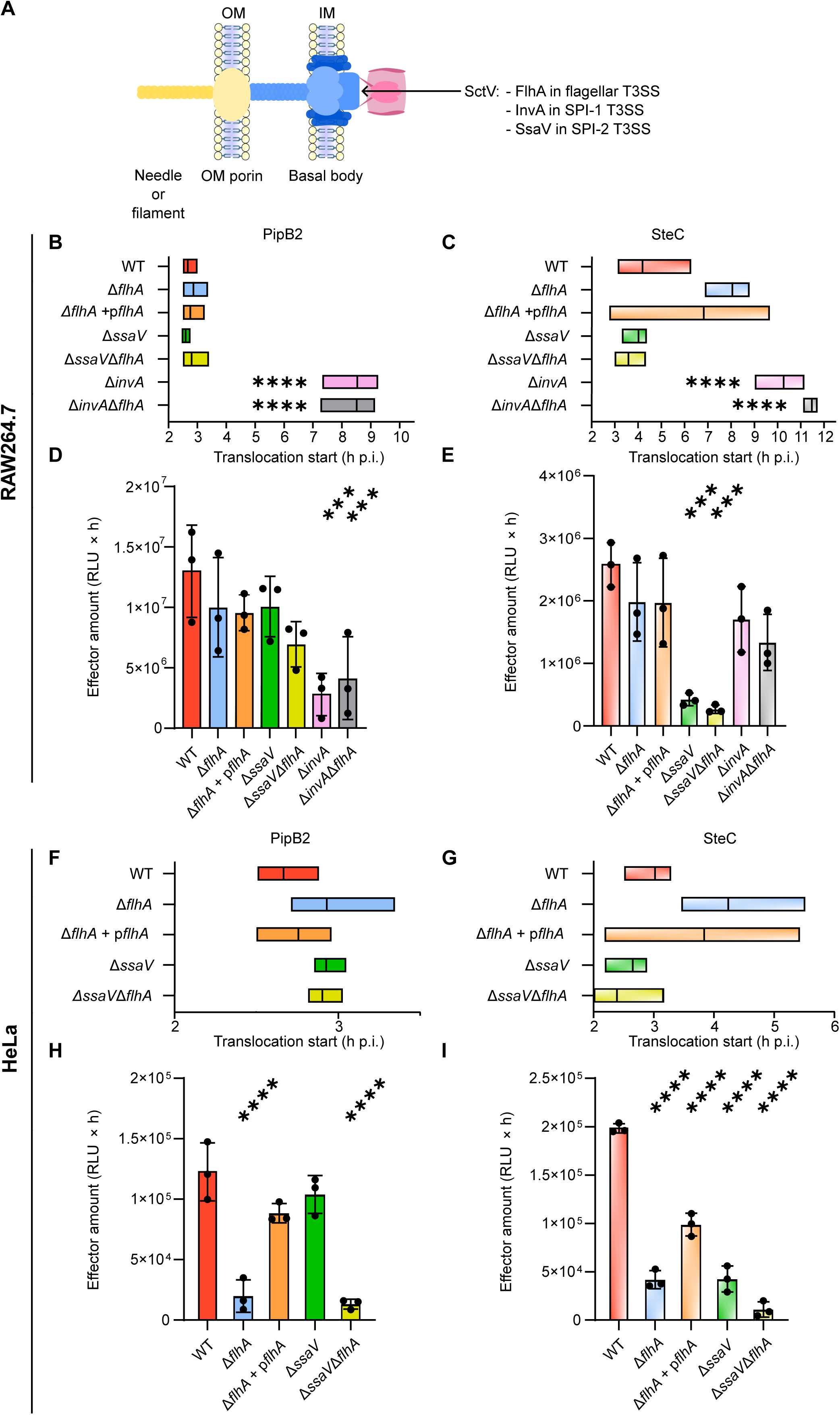
Flagellar T3SS is involved in effector translocation into epithelial cells. **A.** Schematic structure of T3SS with highlighted position of gate protein SctV. **B, C, F, and G.** Start time of PipB2 (B and C) and SteC (F and G) effector translocation by *Salmonella* WT, flagella gate mutant Δ*flhA*, complemented flagella gate mutant Δ*flhA* + p*flhA*, SPI-2 gate mutant Δ*ssaV*, and double mutant Δ*ssaV*Δ*flhA* SPI-1 gate mutant Δ*invA*, and double mutant Δ*invA*Δ*flhA* into RAW264.7-LgBiT (B and F) and HeLa-LgBiT (C and G) cells. Data from three independent experiments performed in duplicates are shown as floating bars with line at mean. *** p < 0.001 (One way ANOVA with Tukey HSD test). **D, E, H, and I.** Amounts of PipB2 and SteC effectors translocated by intracellular bacteria from experiments shown in B, C, F, and G. Normalised and raw data are shown in Extended Data Fig. 3. *** p < 0.001 (One way ANOVA with Tukey HSD test).

In contrast, *Salmonella* Δ*flhA* mutant exhibited a near-complete loss of PipB2-HiBiT translocation and a marked reduction in SteC-HiBiT translocation in HeLa cells (Fig. 3H and I and Extended Data Fig. 3E to H). Complementation of *flhA* deletion restored PipB2 translocation into HeLa-LgBiT, confirming the involvement of the flagellar T3SS in effector translocation (Fig. 3H and Extended Data Fig. 3E and F). Since the complementation of *flhA* deletion led to only partial restoration of SteC-HiBiT translocation (Fig. 3I and Extended Data Fig. 3H), we performed a motility assay using *Salmonella* WT, Δ*flhA*, and Δ*flhA* + p*flhA* strains to examine flagellar function in the complemented strain. As expected, motility was fully restored in the complemented strain (Extended Data Fig. 4), confirming that the complementation is functional. This, however, suggests that the translocation and motility functions of flagella may not overlap completely. Together, these findings also show that flagellar machinery contributes to effector translocation and may coordinate with canonical virulence-associated T3SSs during infection.

### SctV and SctN of flagellar T3SS share common structure with that of injectisomes

To determine whether flagellar T3SS components regulating secretion of flagellar proteins are physically capable of supporting effector translocation, we analysed structural similarity of SctV gate components and of the ATPases SctN powering secretion among the individual T3SS. We performed a comparative protein structure analysis using the FATCAT (Flexible structure Alignment by Chaining AFPs) algorithm and ChimeraX 1.10.1. visualisation. FATCAT alignment of SctV components (flagellar FlhA, SPI-1 injectisome InvA, and SPI-2 injectisome SsaV) as well as of SctN (flagellar FliI, SPI-1 injectisome InvC, and SPI-2 injectisome SsaN) showed high structural similarity between the corresponding components of flagellar T3SS and its injectisome counterparts. ChimeraX superimposed structures are shown in Figure 4 and results of FATCAT analysis are in Extended Data Table 2. Combined alignment of all SctV proteins is presented in Extended Data Movie 1 and of SctN proteins in Extended Data Movie 2. Comparison of primary amino acid sequence of the SctV and SctN components are quantified in Extended Data Table 1 and shown in Extended Data Figure 5 and 6, respectively. Together, this analysis shows that both SctV and SctN components share strong structural conservation among all three *Salmonella* T3SS, consistent with their common role in T3SS-mediated protein translocation.

**Figure 4.**
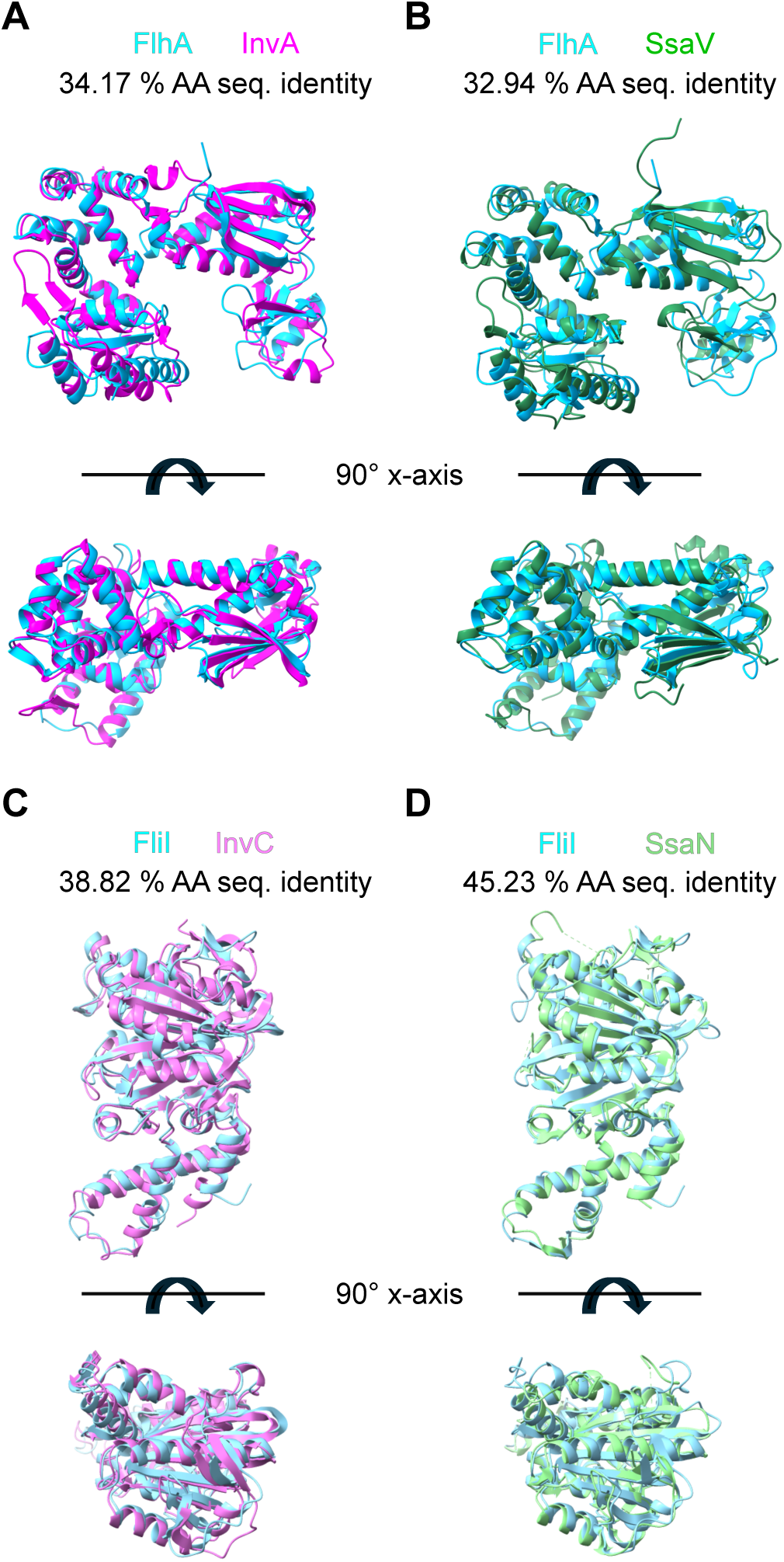
T3SS gate and ATPase proteins share an overall structural fold. **A-B**. Structural alignment of flagellar gate cytosolic domain (FlhA_cyt_) with corresponding proteins from (**A**) SPI-1 (InvA_cyt_) or (**B**) SPI-2 (SsaV_cyt_) injectisomes. **C-D**. Structural alignment of flagellar FliI ATPase with corresponding proteins from (**C**) SPI-1 (InvC) or (**D**) SPI-2 (SsaN) injectisomes. The N terminal domain not visible in InvC and SsaN crystal structures is not shown. PDB IDs used are listed in brackets: InvA (2X49), FlhA (6CH1), SsaV (7AWA), InvC (6RAE), FliI (2DPY; the structure presented starts with K108), and SsaN (4NPH). Figure was prepared using ChimeraX 1.10.1.

### Efficient effector translocation depends on intact flagellar structure

Since the flagellar hook and filament are major structural distinctions between the flagellar T3SS and injectisomes, we analysed the structural differences of the individual protein complexes constituting the channel lumen. Similarly to SctV and SctN, the overall fold is the same in all analysed structures (Fig. 5A). However, while the inner diameter of injectisome needles (PrgI and SsaG) is similar to the inner diameter of the flagellar hook (FlgE), the inner diameter of flagellin filaments (FliC and FljB) were significantly higher (Fig. 5B). This suggests that the inner diameter of the flagellum allows effector translocation. The internal surface charge (Fig. 5C) of flagellar hook (FliC or FljB) is also in the range of the SPI-1 and SPI-2 injectisome needles (PrgI and SsaG, respectively) indicating that effectors that are translocated by both injectisomes do not face charge restrictions within flagella.

**Figure 5.**
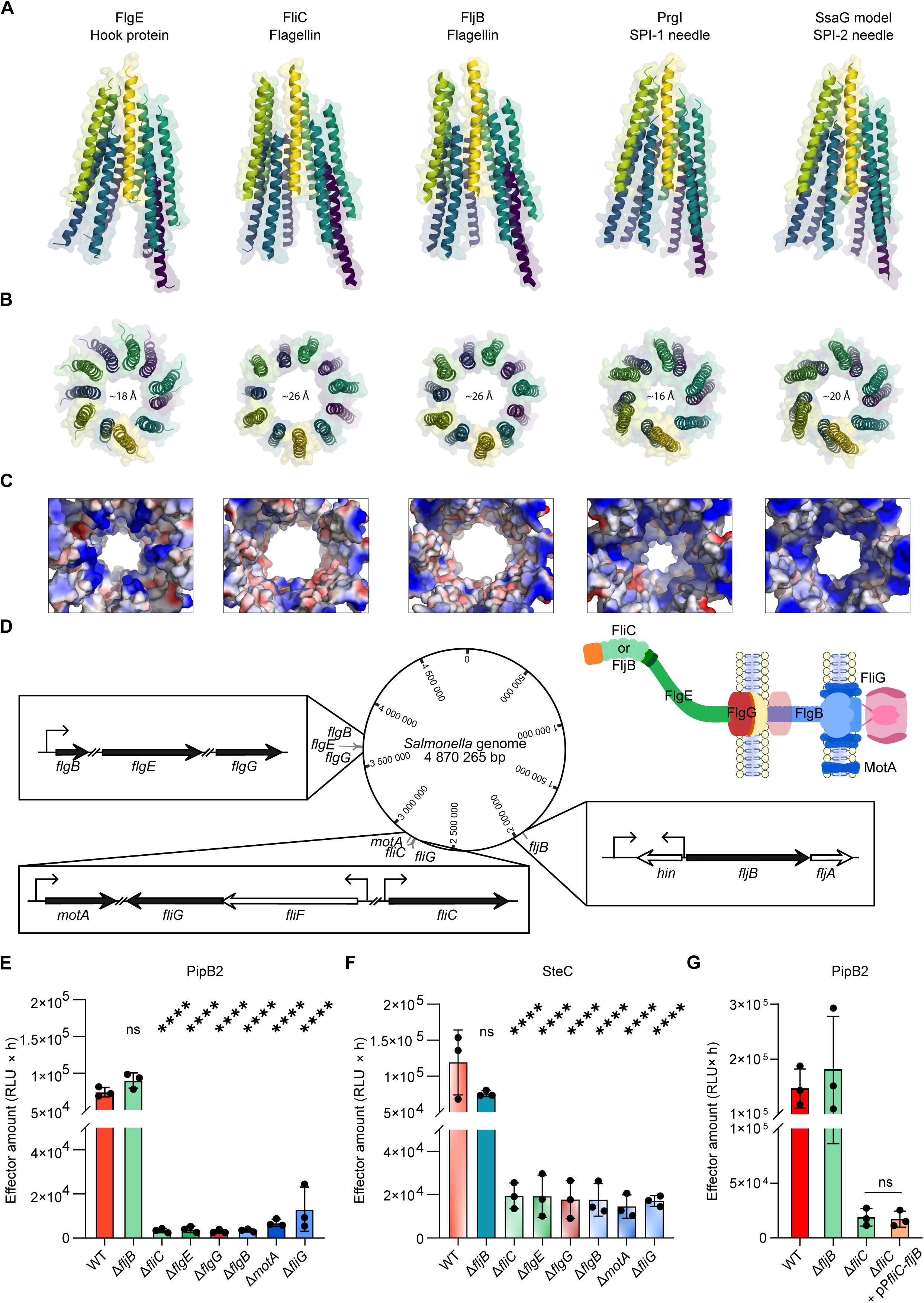
FliC but not FljB flagellar filament is required for efficient effector translocation. **A to C.** Structural comparison of C-terminal regions of flagellar hook (composed of FlgE), filament (FliC or FljB), SPI-1 injectisome needle (PrgI), and SPI-2 injectisome needle (SsaG) that constitute the lumen of each channel. PDB IDs of structures used are listed in brackets: FlgE (6K3I), FliC (9GNZ), FljB (6JY0), and PrgI (6DWB). Model of SsaG was generated using SWISS-MODEL. **D**. Map of *Salmonella* Typhimurium ATCC 14028 chromosome with highlighted position and operon structure of genes used in this figure. Genes mutated in panels B and C are depicted as black arrows, neighbouring genes are depicted as white arrows. Positions of corresponding proteins in the flagellar structure is shown. **E**. **and F.** HeLa-LgBiT cells were infected with *Salmonella* WT, C ring mutant Δ*fliG*, stator mutant Δ*motA*, inner rod mutant Δ*flgB*, outer rod mutant Δ*flgG,* hook mutant Δ*flgE*, and flagellin mutants Δ*fliC* and Δ*fljB*. Amounts of PipB2 (B) and SteC (C) effector translocated by intracellular bacteria from three independent experiments performed in duplicates are shown. Normalised and raw data are shown in Extended Data Fig. 7. ns, not significant; ns, p > 0.05; **** p < 0.0001 (One way ANOVA with Tukey HSD test). **G.** HeLa-LgBiT cells were infected with *Salmonella* WT, Δ*fljB,* Δ*fliC, and* Δ*fliC +* pP*flic-fljB*. Amounts of PipB2 effector translocation by intracellular bacteria from three independent experiments performed in duplicates are shown. Normalised and raw data are shown in Extended Data Fig. 7. ns, p > 0.05 (One way ANOVA with Tukey HSD test).

**Figure 6.**
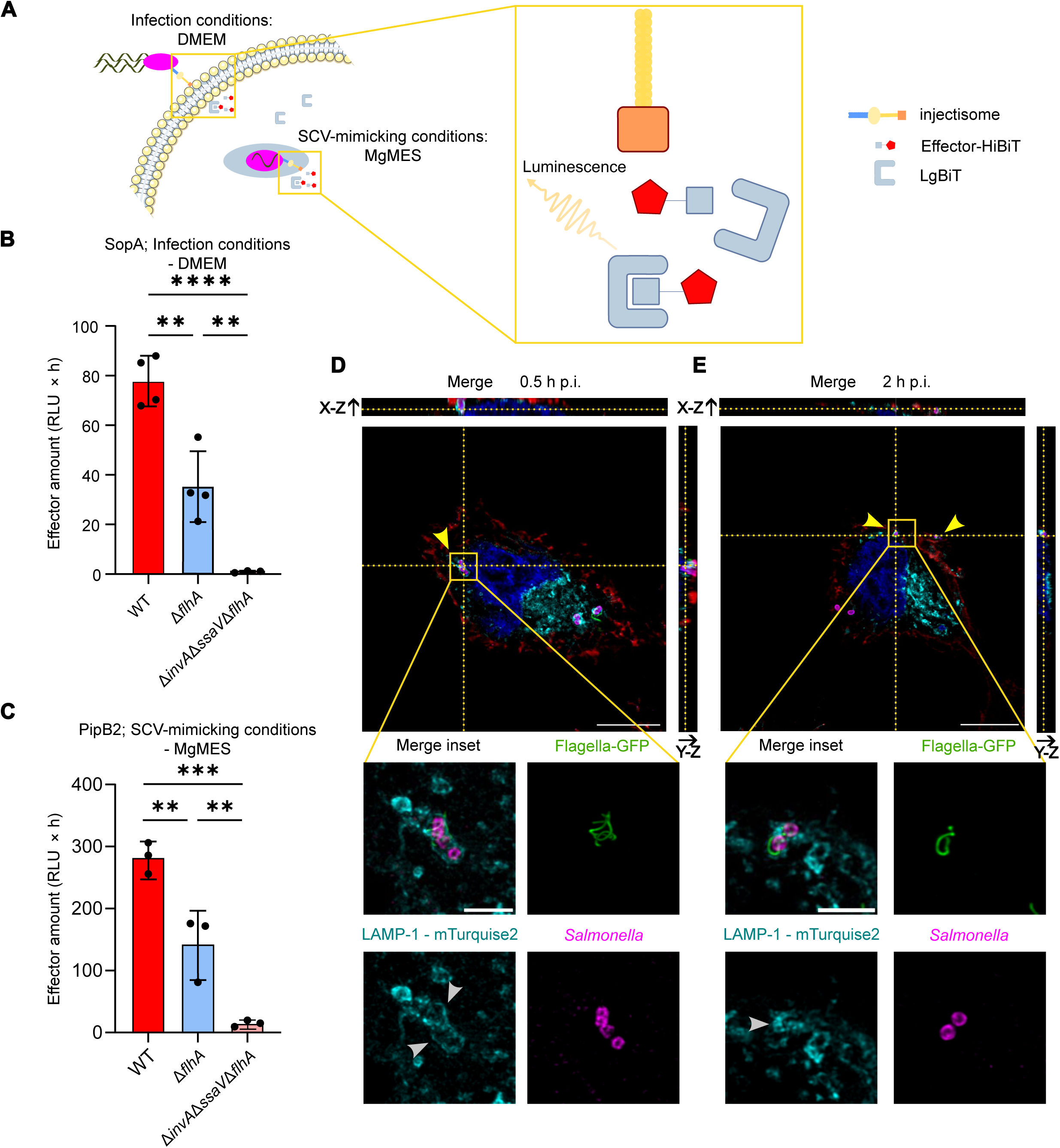
Translocation-proficient flagella are preserved within SCV. **A.** Schematic representation of effector translocation/secretion condition comparison. Secretion of SopA-HiBiT into DMEM (mimicking infection condition) or PipB2-HiBiT into MgMES pH 5.0 (mimicking conditions within the SCV). The medium contained extracellular LgBiT for efficient complementation of Nano-Luc. **B and C.** Secretion of SopA-HiBiT into DMEM (B) used for HeLa cell infection experiments or PipB2 into MgMES pH 5.0 (mimicking conditions within SCV; C). Experiments were normalised by setting the maximum luminescence signal of each repetition to 100%. Data from four or three independent experiments are shown. ** p < 0.01, *** p < 0.001, **** p < 0.0001 (One-way ANOVA with Tukey HSD test). **D and E.** Representative confocal microscopy images depicting localisation of flagella within epithelial cells. HeLa cells expressing SCV marker LAMP-1-mTurquoise2 (cyan) were infected with *Salmonella glmS::*mCherry WT (MOI 1:100; magenta) for 0.5 h p.i. (D) or 2 h p.i. (E). Host cell membranes were labelled with WGA (red), fixed, and permeabilized for labelling of flagella (green) and nuclei (DAPI, blue). The yellow arrowheads point towards intracellular flagellated bacteria. The grey arrowheads points towards SCVs. Scale bar, 5 and 1 µm.

To identify the parts of flagella crucial for effector translocation, we generated *Salmonella* strains with deletion of genes corresponding to different parts of flagellar T3SS: Δ*motA* (motor-stator), Δ*fliG* (rotor), Δ*flgB* (proximal rod), Δ*flgG* (distal rod), Δ*flgE* (hook), or Δ*fliC* and Δ*fljB* (mutually exclusive types of flagellins)^20^ (Fig. 5D). The selected genes are parts of different chromosomal regions^31^ - region 1 (*flgE*, *flgG*, *flgB*), region 2 (*motA*), region 3 (*fliC*, *fliG*), and region H2 (*fljB*) - and are organised in an independent operon structures^32^ which excludes the possibility of polar effects between individual operons (Fig. 5D). Despite their distinct locations, deletion of any of the genes except *fljB* resulted in approximately 10-fold reduction in effector translocation into HeLa-LgBiT cells (Fig. 5E and F). These results indicate that an intact flagellar structure is required for flagella-mediated effector translocation. Importantly, complementation of *fliC* deletion with *fljB* under the control of *fliC* promoter did not restore PipB2-HiBiT translocation (Fig. 5G) suggesting that the flagellar filament composed of FliC but not of FljB is required for effector translocation. Although FljB possess an extra domain in comparison to FliC, this is localised outside of the filament lumen and does not have any obvious impact on filament lumen diameter or charge. Therefore, why the flagellum composed of FljB fails in effector translocation is not clear and remains the focus of our further investigation.

### Flagellum translocates effectors in both SPI-1 and SPI-2 inducing conditions

We observed that flagella-mediated effector translocation is involved in *Salmonella* invasion into epithelial cells (Figs. 1 and 5). To examine if conditions within SCV allow prolonged flagellar effector delivery, we compared *in vitro* flagella-mediated effector secretion into the infection medium (DMEM) and SCV mimicking minimal medium (MgMES pH 5.0) based on the split-luciferase system (Fig. 6A). Deletion of *flhA* leads to a significant decrease of SopA secretion into DMEM in the SPI-1-inducing conditions used for HeLa cell invasion (Fig. 6B). Similarly, PipB2 secretion into the SCV-mimicking minimal medium MgMES pH 5.0 was reduced in the absence of *flhA* (Fig. 6C), suggesting that flagella can translocate effectors into the host cell cytosol from the extracellular space as well as from the SCV.

### *Salmonella* is internalised with FliC flagellar filament

For flagella to support effector translocation from SCV before SPI-2 injectisome induction, flagellar apparatus must be physically present in the SCV. Therefore, we examined the presence of flagella within host cells using confocal microscopy of HeLa cells infected with *Salmonella* WT at 0.5 and 2 h p.i. Flagella were detected on extracellular bacteria at 0.5 h p.i. (Fig. 6D). Importantly, we observed flagella structure on intracellular bacteria surrounded with Lamp1^+^ endosomal membrane at 0.5 and 2 h p.i. (Fig. 6D and E, respectively) but not at 16 h p.i. This suggests that flagella are internalised with *Salmonella* during invasion and are associated with the bacterium within SCV.

Overall, our results indicate that *Salmonella* flagellar structures are similar to those of both SPI-1 and SPI-2 injectisomes. The FliC flagellar filament translocates effector proteins from the extracellular environment supporting *Salmonella* invasion into non-phagocytic cells. Furthermore, the flagella-mediated effector translocation continues from the early SCV until the SPI-2 injectisome is induced. Thus, effector translocation through FliC-composed flagella has an important function in *Salmonella* virulence.

## Discussion

*Salmonella enterica* encodes three distinct T3SS: the SPI-1- and SPI-2-encoded injectisomes and the flagellar T3SS. Here, we demonstrate that the flagellar T3SS supported SPI-1–mediated invasion through translocation of canonical injectisome effector proteins into the host cell cytosol. Flagella secreted effectors into the extracellular milieu and continued to translocate effectors after establishing contact with host cells. Following closure of the host membrane around *Salmonella*, flagella tightly wrapped around the bacterium were retained within the nascent SCV and continued to translocate effectors into the host cell cytosol.

Beyond motility, flagella from several bacterial species, including *Salmonella*, have been implicated in adhesion to host cells^23,33^. Although we observed a mild but statistically non-significant reduction in adhesion of *Salmonella* Δ*flhA* mutant lacking extracellular flagellar structure, invasion into host cells was markedly impaired (Fig. 1). This data indicate that flagella contribute primarily to invasion rather than adhesion, most likely through their role in effector translocation. In addition, SPI-1-independent invasion into several non-phagocytic cell types was reported^34^. Hypothetically, this invasion could be mediated by flagellar effector translocation.

We detected flagella-dependent effector secretion into the extracellular medium (Fig. 6B) and observed effector delivery in minimal medium mimicking the SCV environment (Fig. 6C). These results indicate that flagella-mediated effector translocation can begin immediately upon first host cell contact and persist throughout bacterial internalization and nascent SCV formation, following SPI-1 downregulation and prior to full SPI-2 induction.

Substrate translocation through T3SS is a tightly regulated process initiated by interaction of the substrate with its cognate chaperone. The chaperone interaction drives the selection and specificity of translocation machinery^35,36^ . Following recognition of the substrate by the T3SS machinery, the substrate enters the secretion system through the complex of gate proteins SctV in a process powered by the SctN ATPase. Both SctV and SctN share high degree of sequence and structural similarity between all *Salmonella* T3SS (Figs. 4, Extended Data Fig. 5 and 6, and Extended Data Table 1). The T3SS machinery then ensures transfer of the effectors through both bacterial and host membranes. While absence of the tip complex of injectisomes does not prevent effector secretion^37,38^, the complete structure of flagella including the filament was necessary for effector translocation (Fig. 5).

*Salmonella* flagella filaments can be composed of either FliC or FljB flagellin. Phase variation between these two flagellins is controlled by the Hin recombinase^39^. Switching from FljB to FliC occurs more frequently than in the opposite direction, resulting in preferential FliC expression in the population^40^. We show that deletion of *fljB* did not affect effector translocation (Fig. 5B and C), despite detectable *fljB* expression under the experimental conditions (Fig. 2G). In contrast, deletion of *fliC* completely abolished flagella-mediated effector translocation (Fig. 5B and C). This observation is consistent with the reported importance of FliC during *in vivo* infection. Although *fljB* mutants are attenuated in a mouse infection model^41^, FljB-composed filament is not sufficient for full invasion into epithelial cells^42^. In contrast, FliC alone appears sufficient for invasiveness in people, as the invasive nontyphoidal *Salmonella* ST313 lineage II.1 evolving in sub-Saharan Africa has lost *fljB*^43^. Together, these findings suggest that FliC-mediated effector translocation contributes to *Salmonella* virulence during natural infection.

In contrast to our observation (Fig. 2), flagellar genes are downregulated more rapidly than SPI-1 genes following entry into HeLa cells^44^, and proteomic analysis did not detect a substantial decrease in SPI-1 injectisome-associated proteins between 1 and 6 h p.i. in HeLa cells^45^. However, a significant proportion of intracellular *Salmonella* in HeLa cells may reside in the cytosol and re-induce SPI-1 expression by 6 h p.i.^46–48^. Thus, the apparent stability of SPI-1 protein levels may reflect averaging across distinct bacterial subpopulations providing a possible explanation for an apparent discrepancy with our observations, which showed persistent flagellation and rapid drop of SPI-1 T3SS mediated translocation.

During T3SS-mediated translocation, substrates pass through a narrow central channel^49,50^ within the secretion apparatus. Several *Salmonella* effectors are translocated by both SPI-1 and SPI-2 injectisomes^30^, indicating that both these structures accommodate effector transit in terms of channel diameter and electrostatic properties. Notably, both the diameter and inner surface charge of the flagellar hook and filament fall within the range of those of the SPI-1 and SPI-2 injectisomes (Fig. 5B and C), suggesting that physical constraints are unlikely to limit effector passage through the flagellar filament.

We detected intact flagella within the SCV after the invasion and at 2 h p.i. (Fig. 6E), indicating that the invasion process can enclose and internalize large portion of flagellum into the SCV, thus supporting a role for flagella-mediated effector translocation during early intracellular stages. However, flagella were no longer detectable in SCV late during infection, suggesting that changes in SCV composition between 2 and 16 h p.i. promote flagellar disassembly. During this interval, SCV undergoes progressive acidification to pH of approximately 4.5–5.5^51,52^. Although acidification below pH 5.0 induces structural changes in flagella without causing disassembly^53^ complete filament dissociation requires pH 2.3^54^, a condition unlikely to occur within the SCV. These observations suggest that additional host or bacterial factors contribute to flagellar disassembly in the vacuolar environment.

We did not identify the specific flagellar component responsible for pore formation in host membranes. However, it is possible that no dedicated pore-forming structure is required, and that membrane penetration occurs during enclosure of the nascent SCV. The flagellin filament itself would then traverse the host membrane in the place where membrane ruffles were not able to cover the whole flagellum. Consistent with this possibility, electron microscopy has shown that *Escherichia coli* flagella can penetrate host plasma membrane in a FliC- and motility-dependent manner^55^.

In summary, we demonstrate that fully assembled flagella mediate translocation of *Salmonella* effectors both from the extracellular space and from the SCV. This activity contributes to the invasion of non-phagocytic cells and persists until SPI-2 injectisome induction. Our findings expand the established hierarchy of *Salmonella* effector translocation. Upon host cell contact, early effectors are delivered via both the SPI-1 injectisome and flagella. Following internalization, late SPI-1 injectisome effectors continue to be translocated in part by flagella. Subsequently, early SPI-2 injectisome effectors are initially delivered through flagella before SPI-2 injectisome-dependent secretion predominates, culminating in late SPI-2 effectors that are exclusively translocated via the SPI-2 injectisome. This model presents a mechanism of continuous translocation of *Salmonella* T3SS effectors and shows important function of flagella in creation of *Salmonella*-permissive niche within host cells. Flagella-mediated effector translocation, therefore, represents an important and generalizable virulence mechanism.

## Methods

### Bacterial strains

Bacteria were grown in Luria–Bertani (LB) medium or in SPI-2-inducing medium MgMES (170 mM 2-(N-morpholino)ethanesulfonic acid (MES) at pH 5.0, 5 mM KCl, 7.5 mM (NH_4_)_2_SO_4_, 0.5 mM K_2_SO_4_, 1 mM KH_2_PO_4_, 8 mM MgCl_2_, 38 mM glycerol, and 0.1% casamino acids), supplemented with ampicillin (100 µg/ml), kanamycin (70 µg/ml), or chloramphenicol (30 µg/ml), as appropriate. Gene deletion strains were constructed using the λ-red recombination system by replacing the targeted gene with an antibiotic resistance cassette^56^. See Extended Data Table 3 for all bacterial strains.

### Plasmid construction

The effector protein genes under their native promoters were inserted into the pWSK29 plasmid using XhoI - BamHI / HindIII - BamHI. A complementation plasmid for *ΔflhA* deletion was constructed using the pET-28b backbone to introduce *flhBAE* operon between NcoI-XhoI cloning sites.

To stably express GFP from the bacterial chromosome, GFP under the control of P*trc* was inserted into the chromosomal Tn7 loci using pGRG25. P*trc*-GFP was cloned into the pGRG25^57^ using NotI - SphI and subsequently integrated downstream of *glmS* on the chromosome, creating *S*Tm-GFP strains. Similarly, *Salmonella* strain expressing mCherry (*S*Tm-mCherry) was constructed between NotI - SphI restriction sites as described in Pospíšilová et all^58^. All primers are listed in Extended Data Table 4, and all plasmids are listed in Extended Data Table 5. All plasmids were verified by Sanger sequencing.

### Cell Culture

Human HeLa LgBiT, a kind gift from Jana Kamanova (Infection Biology Laboratory, Institute of Microbiology of the Czech Academy of Sciences^59^) and RAW264.7 mouse macrophages were maintained in Dulbecco’s modified eagle medium (DMEM; Sigma) supplemented with 10% heat-inactivated foetal calf serum (FCS; Sigma).

### Transfection and virus production

For transient plasmid transfections, pLamp1-mTurquoise2 (Addgene #98828) and Lipofectamine 2000 (Thermo Fisher Scientific) were combined according to the manufacturer’s instructions and incubated in OptiMEM for 5 min at room temperature before being added to cells. Cells were infected 16–20 h after plasmid transfection.

RAW264.7 stably expressing LgBiT were prepared using lentiviral transduction. VSV-pseudotyped viruses were prepared by co-transfection of 250 ng pLX304_CMV::mCherry-LgBiT-FLAG-NES (Addgene #199715), 100 ng pCMV-VSV-G (Addgene #8454), and 150 ng psPAX2 (Addgene #12260) into HEK293T cells seeded in a 24-well plate using Lipofectamine 2000 (Thermo Fisher Scientific) as described above. The culture medium was replaced 24 h after transfection and the supernatant containing viruses was collected 48 h after transfection and filtered through a 40 µm (Avantor) filter. Lentiviruses were added to RAW264.7 together with polybrene (Sigma) at 10 μg/ml. At 24 h post-transduction, cells were sorted by flow cytometry for the mCherry fluorescence.

### Cell Infection

HeLa cells and RAW264.7 macrophages were infected with a subculture of *S.* Typhimurium at the late exponential phase (OD_600_ = 1.8) with an MOI of 100:1 or 20:1, respectively, for 30 min. To induce cell contact for non-motile strains, samples were centrifuged at 100 × *g* for 5 min. Cells were washed 3 times with PBS and incubated in fresh medium containing gentamicin (100 µg/ml) to kill extracellular bacteria. After 1 h, the antibiotic concentration was reduced to 20 µg/ml, and cells were processed at various time points post-infection as indicated in the figure captions.

### Split luciferase assay

To assess the translocation of secreted effectors, we measured the luminescence generated by the reconstitution of split NanoLuc luciferase. LgBiT was either expressed in host cells or added externally into the culture medium, while HiBiT was fused to the translocated effector, and luminescence was recorded following infection as described above.

For HeLa cells, the Nano-Glo Live Cell Assay System (Promega) was used, whereas RAW264.7 macrophages were assayed with Endurazine Live Cell Substrate (Promega). Fully reconstituted Nano-Glo Live Cell Reagent was prepared by combining 1 volume of Nano-Glo Live Cell Substrate with 19 volumes of Nano-Glo LCS Dilution Buffer (20-fold dilution). For each well, 25 μL of reagent was added to 100 μL of phenol red–free DMEM (Gibco) supplemented with 5% heat-inactivated foetal calf serum (FCS; Gibco). Endurazine Live Cell Substrate was diluted 100-fold in 200 μL of phenol red–free DMEM containing 5% heat-inactivated FCS (Sigma). Both substrates are based on furimazine; Endurazine is covalently modified with a protective group (R) to enable prolonged luminescence detection.

Extracellular detection of translocated effectors was done using Nano-Glo HiBiT Extracellular Detection System (Promega). This set-up includes dilution of the LgBiT Protein in a ratio of 1:200 and the Nano-Glo® HiBiT Extracellular Substrate in a ratio of 1:25 with an appropriate volume of Nano-Glo® HiBiT Extracellular Buffer. For each well, 100 μL of prepared reagent was mixed with an equal amount of phenol red–free DMEM (Gibco) supplemented with 5% heat-inactivated foetal calf serum (FCS; Gibco). All luminescence measurements were performed in white 96-well plates with transparent bottoms (Corning) using a Spark® multimode microplate reader (Tecan), following the acquisition settings recommended by the manufacturers.

Detection of the beginning of translocation as shown in Figs. 2 and 3 was calculated using the custom Python script (Appendix A). Onset times were determined by fitting an asymmetric Gompertz model to the rising phase of each effectors’ translocation pattern.

### Fluorescence microscopy

To analyse flagellation within host cells, HeLa cells expressing LAMP-1-mTurquoise2 seeded on glass coverslips were infected with *S*Tm-mCherry as described above. At 0.5, 2, and 18 h p.i. cells were washed with PBS and labelled live with 5 µg/ml of WGA for 10 minutes on ice, washed with PBS and fixed in 4 % paraformaldehyde in PBS at 37°C for 20 min, following labelling with 0.5 µg/ml DAPI for 5 minutes. All samples were permeabilised using 0.2% Triton X-100 for 5 min. and incubated with 2 µg/ml Anti-Flagellin FliC (Invitrogen) monoclonal antibody for 1 h, washed, and incubated with 4 µg/ml of secondary goat anti-mouse IgG polyclonal antibodies (Thermo Fisher Scientific, cat. n. AB_2534069) for 1 h. For further visualisation, coverslips were mounted onto glass slides using Aqua-Poly/Mount (Polysciences) and imaged using a Stellaris 8 inverted confocal laser-scanning microscope (Leica Microsystems, Germany).

A complete list of all antibodies used in the study is presented in Extended Data Table 6.

### Image analysis

All image processing steps were performed using ImageJ/ Fiji and LAS X software.

### Flow cytometry

The ability of bacterial strains to efficiently adhere and invade host cells was assessed by flow cytometry. HeLa cells were infected as described above for 0.5 and 2 h with *Salmonella* strains stably expressing GFP. To detect bacteria that were adherent to the host but not invaded, samples were incubated with polyclonal 1 µg/ml of CSA-1 anti-*Salmonella* antibody (Avantor) and 4 µg/ml of polyclonal donkey anti-Goat IgG antibody (Thermo Fisher Scientific) for 1 h each with 3 washing steps in between. To analyse the expression of fluorescent signals in individual bacteria, the infected host cells were lysed by osmotic lysis using dH_2_O. Freshly prepared samples were processed on the same day using ZE5 (Biorad) cytometer. Data was analysed using FlowJo v10 (BD) software.

### Motility assay

Motility of individual *S.* Typhimurium strains was tested on 0.5% agar plates containing LB. The agar was spiked with indicated *S.* Typhimurium strains at the late exponential growth phase (OD_600_ = 1.8) and incubated at 37°C. The motility was evaluated by measuring the total square of the bacterial swimming zone from the point of inoculation after 18 h of incubation. The experimental setup is based on a modified adaptation of the methodologies previously described by Soria-Bustos et al. and Partridge and Harshey^60,61^. Calculation of the swimming area for the motility assay was done using the “Freehand Selections” tool and automatically measured in ImageJ as described in Morales-Soto et al.^62^

### Colony-forming unit counting assays

Rates of intracellular replication were determined by the colony-forming unit count. HeLa cells infected with *S.* Typhimurium were osmotically lysed at 0.5, 2, and 18 h p.i. Fold replication values were assessed by dividing the CFU count at 18 h p.i. by the CFU count at 2 h p.i. from the respective samples.

CFU/mL = (Number of colonies) × (Dilution factor) / (Volume plated in mL)

### Western blot

To validate the expression and translocation of effectors by Western blot, HeLa cells infected with *Salmonella* were lysed at different time points in lysis buffer (Tris-HCl pH = 8, 220 mM, NaCl 100 mM, EDTA 10 mM, 1 mini tablet of protease inhibitor cocktail, 0.1% Triton) at 4°C for 10 min, and the post-nuclear supernatant was collected after centrifugation at 16,000 × *g* at 4°C for 10 min and analysed by SDS-PAGE and immunoblotting. Bacterial samples used to check the secretion deficiency of mutant strains were lysed with a specialised buffer (1% Triton, 20 mM Tris-HCl, pH = 8.2, 100 mM NaCl, 10 mM EDTA) supplemented with Roche Complete protease inhibitors. This was followed by 30s water-bath sonication (Tesla). Effector proteins were detected using Anti-HiBiT Monoclonal Antibody (Promega), HSP70/DnaK Antibody clone 8E2/2 (Novus Biologicals). Anti-PrgJ and Anti-DnaK were used to detect bacterial proteins. Anti-beta Actin antibody (mAbcam 8226) was used as a loading control. As a secondary antibody, an HRP-linked Anti-mouse IgG or Anti-rabbit IgG antibodies were used for signal visualisation using ECL detection reagent SuperSignal West Femto Substrate (Thermo Scientific) in G:Box Chemi XRQ.

### RT-qPCR

Correlation of effectors abundance and expression to the translocation patterns of the luciferase assay was performed by RT-qPCR. After RNA extraction using Phenol-Chloroform from infected cells, the remaining DNA was removed using TURBO DNase^TM^ (Invitrogen). cDNA was prepared using Maxima H Minus Reverse Transcriptase (Thermo Scientific) kit, and 5x diluted cDNA was used for qPCR using EvaGreen PCR master mix (Solis BioDyne). Data are represented as relative amounts of mRNA normalised to housekeeping genes *gapA* and *tufA*, and *ssaG* expression at 1.5 h p. i. of each protein by the delta-delta Ct method. All primers (https://www.ncbi.nlm.nih.gov/tools/primer-blast/) are listed in Extended Data Table 4.

### Protein structure prediction and visualization

Crystal structures of used proteins were obtained from PDB database and compared based on structural similarities using FATCAT^63^. Representative pictures and the movie of superimposed structures were made in ChimeraX 1.10.1.

Structural figures were generated using PyMOL software v2.6.2. Model of SsaG in Figure 5E-G was obtained using SWISS-MODEL^64^ and PrgI structure as template.

### Quantification and statistical analysis

Statistical significance was calculated using one-way or two-way ANOVA analysis with Tukey’s correction and one-sample Wilcoxon test. All statistical analysis was carried out in the GraphPad Prism v9 software.

## Supporting information

Extended Data Movie 1. Structural comparison of gate proteins FlhA, InvA, and SsaV.

Extended Data Movie 2. Structural comparison of ATPase proteins FliI, InvC, and SsaN.

Extended Data Figure 1. Schematic representation of T3SSs structure.

Extended Data Figure 2. Hierarchy of effector translocation.

Extended Data Figure 3. Involvement of flagella in effector translocation.

Extended Data Figure 4. Complementation of flhA deletion fully restore Salmonella motility.

Extended Data Figure 5. Comparison of primary amino acid sequence of the SctV components of flagellar T3SS, SPI-1 and SPI-2 injectisomes.

Extended Data Figure 6. Comparison of primary amino acid sequence of the SctN components of flagellar T3SS, SPI-1 and SPI-2 injectisomes.

Extended Data Figure 7. Impact of deletion of flagellar components on PipB2 and SteC translocation.

Extended Data Table 1. Universal nomenclature of relevant injectisome and flagellar proteins.

Extended Data Table 2. Results of FATCAT analysis of SctV and SctN similarities

Extended Data Table 3. Bacterial strains used in this study.

Extended Data Table 4. Primers used in this study.

Extended Data Table 5. Plasmids used in this study.

Extended Data Table 6. Antibodies used in this study.

Appendix A. Code for detection of effector translocation start from raw luminescent data.

## Data availability

All data are in the manuscript and/or extended data files and are also available on Zenodo (DOI: 10.5281/zenodo.20524674). Further information and requests for raw data or materials should be directed to the corresponding author. Source data are provided with this paper.

## Code availability

The complete Python script used for detection of the beginning of effector translocation in Figs. 2 and 3 can be found in Appendix A.

## Ethics declaration

Competing interests - The authors declare no competing interests.

## Acknowledgments

We thank all members of Laboratory of Bacterial Virulence and Laboratory of Infection Biology of the Institute of Microbiology of the Czech Academy of Sciences for constructive discussions. HeLa-LgBiT cell line was a gift from Jana Kamanova (Institute of Microbiology of the Czech Academy of Sciences). pLX304_CMV::mCherry-LgBiT-FLAG-NES was a gift from Alice Ting (Addgene plasmid # 199715; http://n2t.net/addgene:199715; RRID:Addgene_199715). We thank Zaneta Slavickova and Jan Svoboda from the Flow cytometry for help with flow cytometry. We acknowledge the Light Microscopy Core Facility, IMG, Prague, Czech Republic, supported by grants “National Infrastructure for Biological and Medical Imaging” (MEYS – LM2023050), “Modernization of the national infrastructure for biological and medical imaging Czech-BioImaging” (MEYS – CZ.02.1.01/0.0/0.0/18_046/0016045) and „Modernization of the VVI Czech-BioImaging” (MEYS – CZ.02.01.01/00/23_015/0008205), for their support with the confocal imaging presented herein. Instrumental support was provided by EATRIS-CZ (code 90253), funded by the Ministry of Education, Youth and Sports of the Czech Republic under the project No. LM2023053. We acknowledge support from Talking microbes -understanding microbial interactions within One Health framework (CZ.02.01.01/00/22_008/0004597).

This work has been funded by a grant from the Programme Johannes Amos Comenius under the Ministry of Education, Youth and Sports of the Czech Republic (project Nr CZ.02.01.01/00/22_008/0004597) and by grants from Czech Science Foundation (22-05356S and 24-11259M).

## Extended Data figures

**Extended Data Figure 1. Schematic representation of T3SSs structure.** The cartoon represents a comparison of *Salmonella* T3SS: SPI-1 injectisome (left), SPI-2 injectisome (centre), and flagellar T3SS (right). Components of cytoplasmic platform are shaded in pink, basal body in blue, outer membrane porin and rod in light yellow, needle of injectisomes in dark yellow, translocon proteins of injectisomes and cap protein of flagellar T3SS filament in orange, unique flagellar elements in green. Homologues structures share the same colour coding. Adapted and modified from Diepold et al.^19^, Wisner et al.^20^ and Mondino et al.^65^

**Extended Data Figure 2. Hierarchy of effector translocation. A.** Translocation of SopA is detectable from the beginning of measurement. HeLa-LgBiT (left) or RAW264.7.-LgBiT (right) cells were infected as in Fig. 2. Measurement of SopA-HiBiT translocation was initiated immediately after removal of extracellular bacteria with PBS washing.

**B and C.** Translocation of individual *Salmonella* effectors to epithelial cells over time. HeLa-LgBiT cells were infected as in Fig. 2. The data are shown as normalised to the maximal luminescent signal for each effector (B) or as raw luminescent values (C). Results of one independent experiment are shown.

**D and E.** Translocation of individual *Salmonella* effectors to macrophages in time. RAW264.7-LgBiT cells. The data from one representative experiment in Fig. 2 are shown as normalised to the maximal luminescent signal for each effector (B) or as raw luminescent values (C).

**F**. Expression profiles of effector proteins in HeLa cells infected with *Salmonella* WT for 1.5, 3, 5, 8, and 24. Data were normalised to the mRNA levels of *ssaG* at 1.5 h p.i. for each effector. Expression of *ssaG* was used to allow comparison between individual genes. Values represent the mean of at least three independent experiments performed in duplicate.

**Extended Data Figure 3. Involvement of flagella in effector translocation.** Translocation of effectors PipB2-HiBiT (**A, B, E,** and **F**) or SteC-HiBiT (**C, D, G,** and **H**) to RAW264.7-LgBiT (**A** to **D**) and HeLa-LgBiT (**E** to **H**) over time. The data from one representative experiment in Fig. 3 are shown as normalised to the maximal luminescent signal for each effector (**A, C, E,** and **G**) or as raw luminescent values (**B, D, F,** and **H**).

**Extended Data Figure 4. Complementation of flhA deletion fully restore Salmonella motility.**

**A.** A representative result of motility assay showing swimming zones of *Salmonella* WT, *Salmonella* Δ*flhA*, and *Salmonella* Δ*flhA* + p*flhA*.

**B.** Quantification of the swimming zone area from motility assay. Data are shown as mean values from three independent experiments in technical duplicates. ** p < 0.01 (One-way ANOVA with Tukey HSD test).

**Extended Data Figure 5. Comparison of primary amino acid sequence of the SctV components of flagellar T3SS, SPI-1 and SPI-2 injectisomes.** The identical amino acid residues are highlighted in **red;** the similar amino acid residues are highlighted in yellow.

**Extended Data Figure 6. Comparison of primary amino acid sequence of the SctN components of flagellar T3SS, SPI-1 and SPI-2 injectisomes.** The identical amino acid residues are highlighted in **red;** the similar amino acid residues are highlighted in yellow.

**Extended Data Figure 7. Impact of deletion of flagellar components on PipB2 and SteC translocation.** Translocation of PipB2-HiBiT (**A** and **B**) and SteC-HiBiT (**C** and **D**) effectors to HeLa-LgBiT cells in time. The data from one representative experiment in Fig. 5 are shown as normalised to the maximal luminescent signal for each effector (**A** and **C**) or as raw luminescent values (**B** and **D**).

**Extended Data Movie 1. Structural comparison of gate proteins FlhA, InvA, and SsaV.** The movie was created using the structures presented in Fig. 4. - gate cytosolic domains of FlhA (PDB: 6CH1; cyan), InvA (PDB: 2X49; magenta), SsaV (PDB: 7AWA; green).

**Extended Data Movie 2. Structural comparison of ATPase proteins FliI, InvC, and SsaN.** The movie was created using the structures presented in Fig. 4. - gate cytosolic domains of FliI (PDB: 2DPY; cyan), InvC (PDB: 6RAE; magenta), SsaN (PDB: 4NPH; green).

**Extended Data Table 1. Universal nomenclature of relevant injectisome and flagellar proteins.**

**Extended Data Table 2. Results of FATCAT analysis of SctV and SctN similarities**

**Extended Data Table 3. Bacterial strains used in this study.**

**Extended Data Table 4. Primers used in this study.**

**Extended Data Table 5. Plasmids used in this study.**

**Extended Data Table 6. Antibodies used in this study.**

**Appendix A. Code for detection of effector translocation start from raw luminescent data.**

