## Supplementary figures and images for "Flagellar system translocates T3SS effectors critical for *Salmonella* infection"

### Extended Data Figure 1. Schematic representation of T3SSs structure.

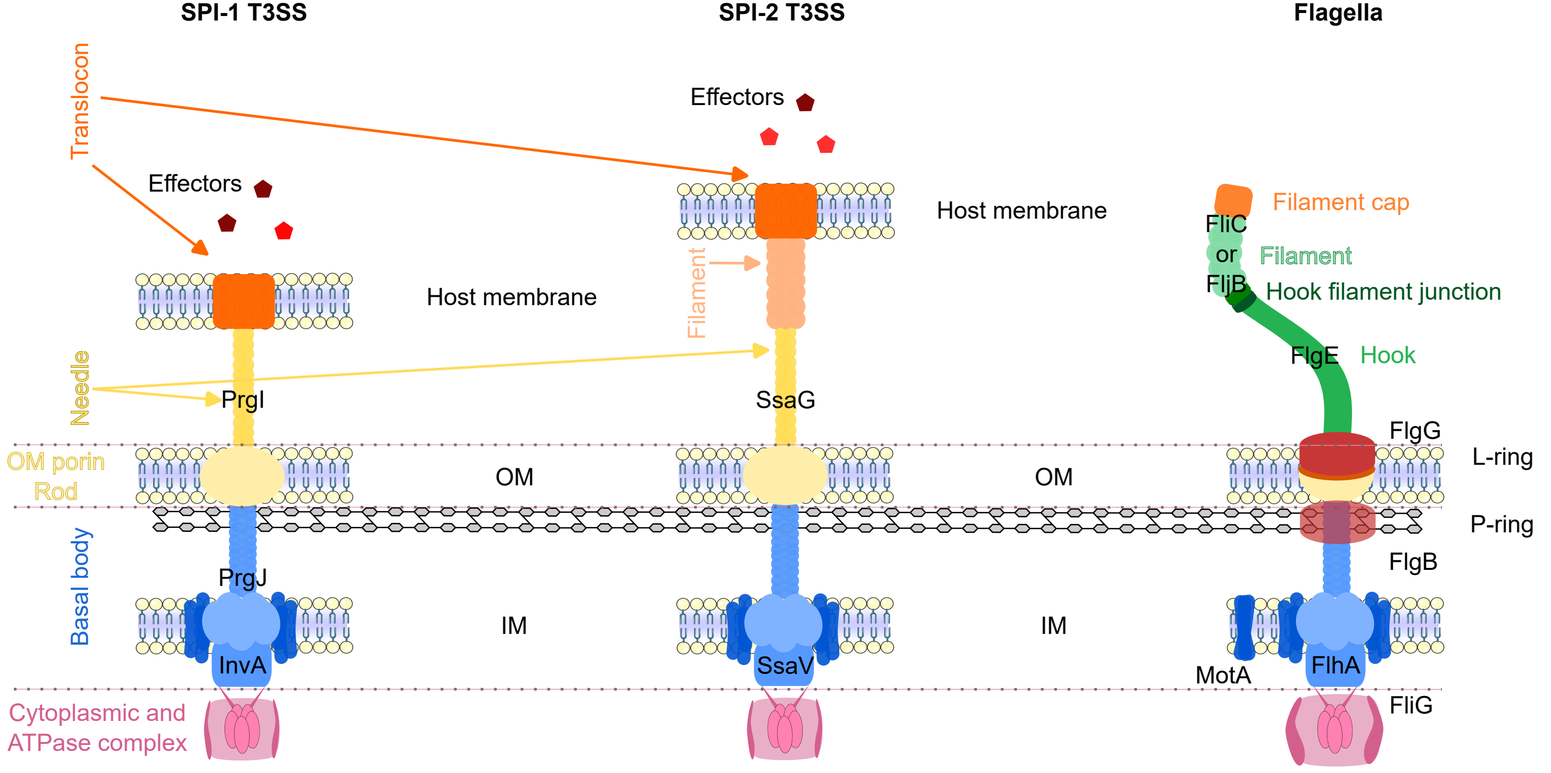

### Extended Data Figure 2. Hierarchy of effector translocation.

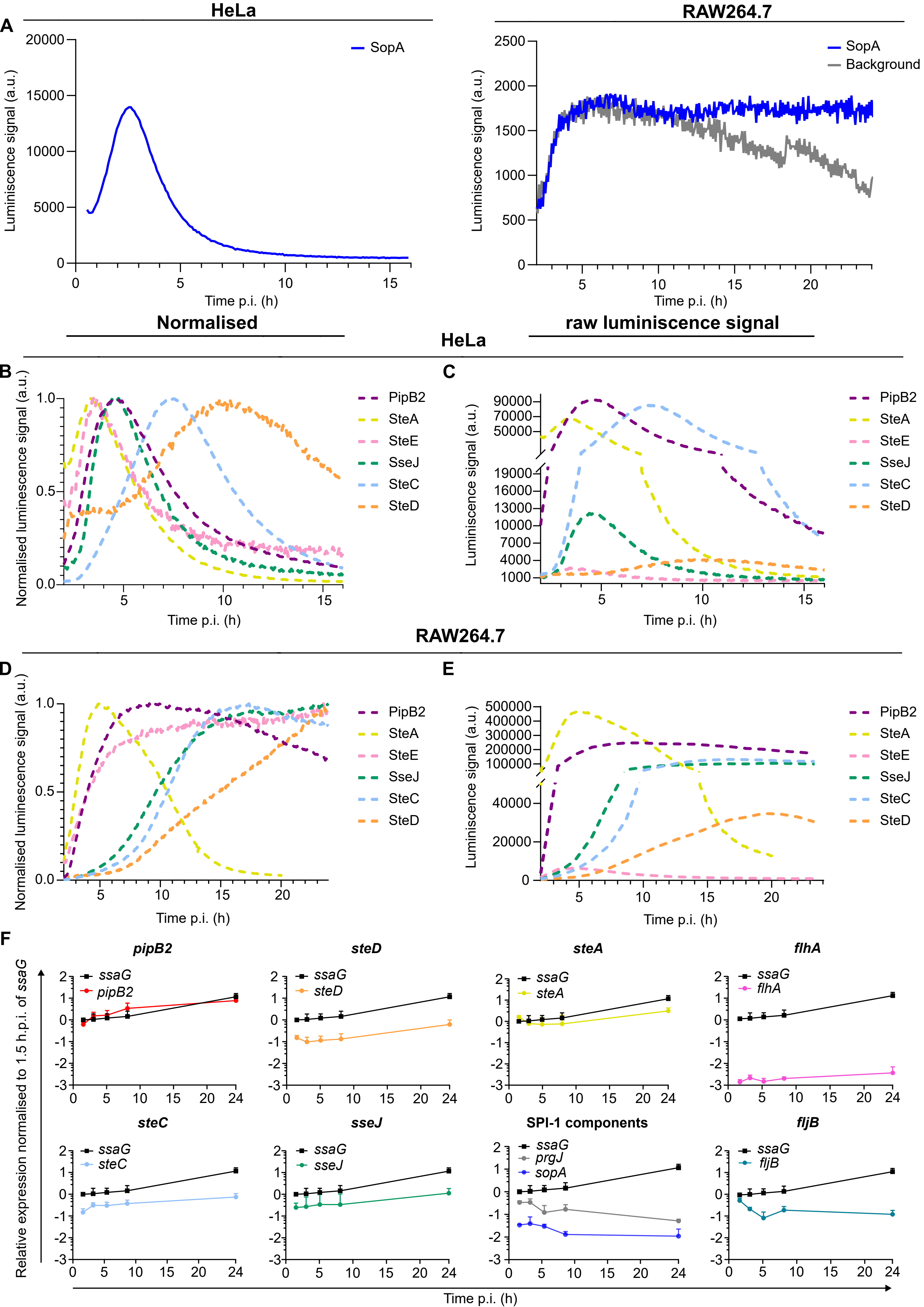

### Extended Data Figure 3. Involvement of flagella in effector translocation.

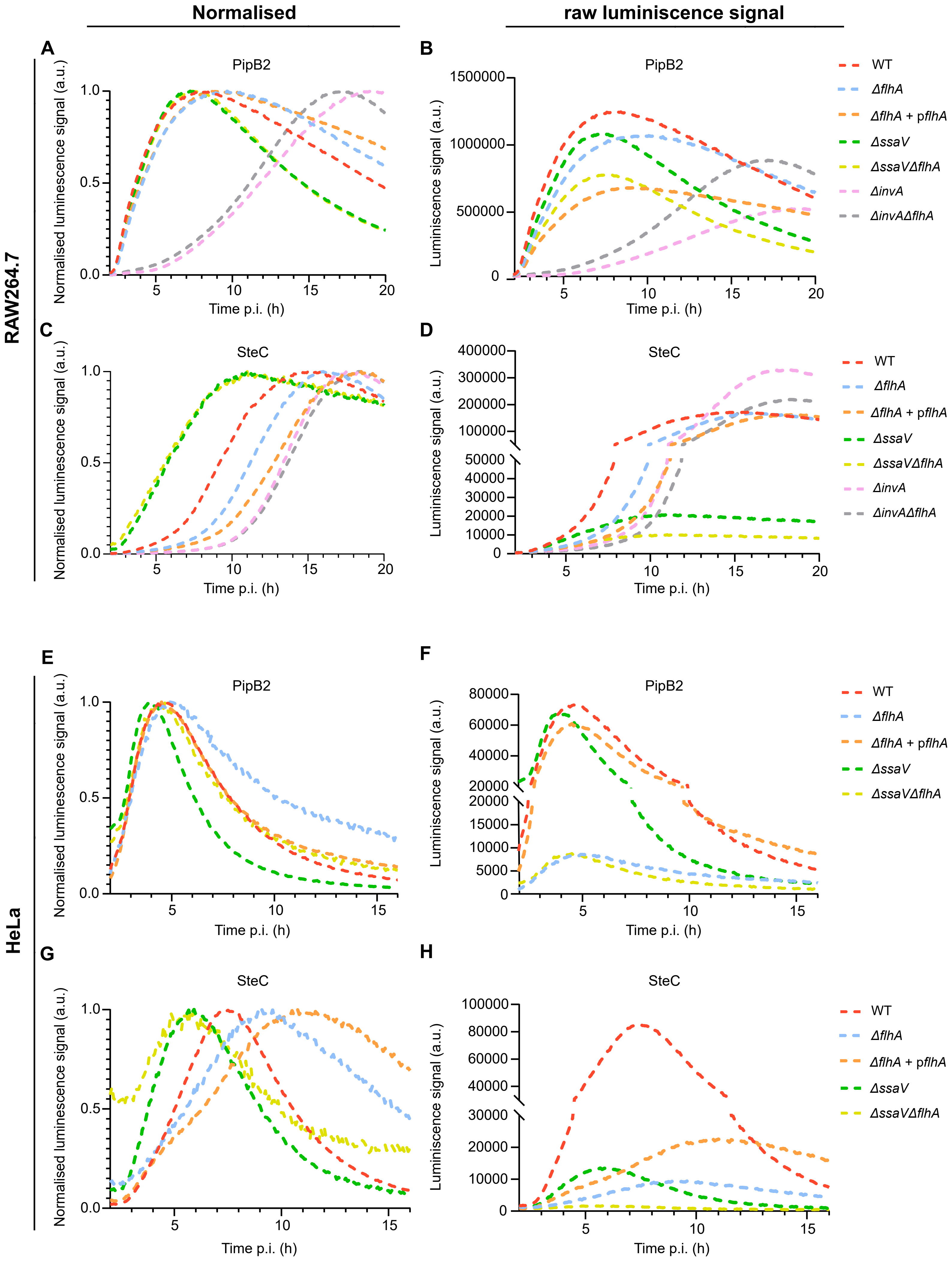

### Extended Data Figure 4. Complementation of flhA deletion fully restore Salmonella motility.

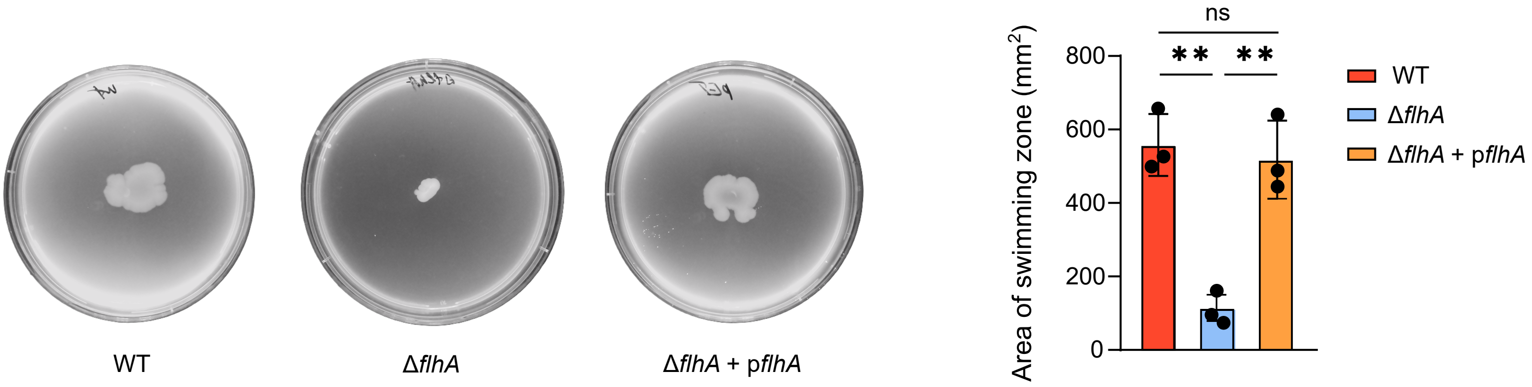

### Extended Data Figure 7. Impact of deletion of flagellar components on PipB2 and SteC translocation.

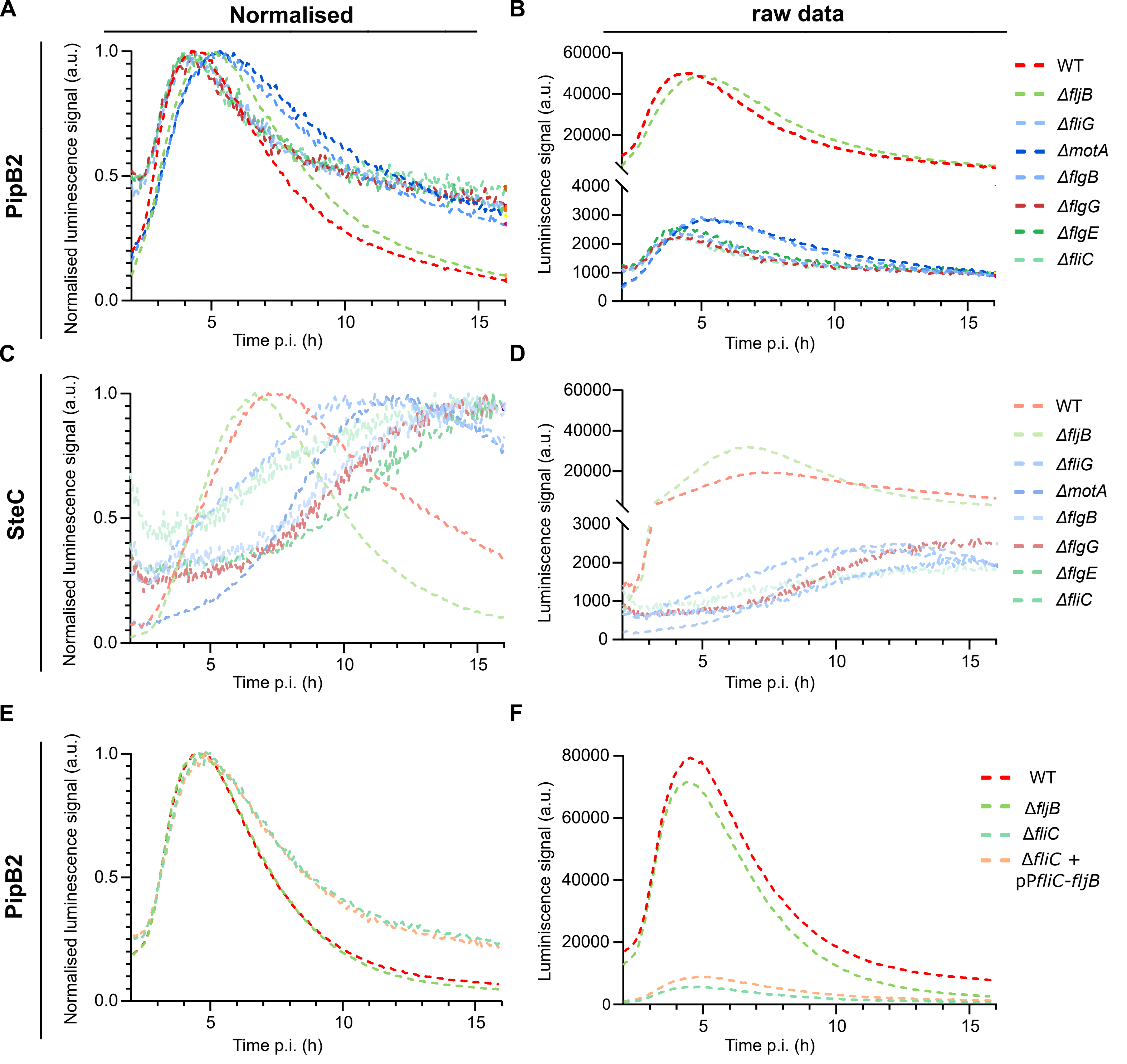
