## Extended Data Figure 5. Comparison of primary amino acid sequence of the SctV components of flagellar T3SS, SPI-1 and SPI-2 injectisomes. for "Flagellar system translocates T3SS effectors critical for *Salmonella* infection"

```
     1       10        20                30        40        50        60
flhA MANLVAMLRLPSNLKSTQWQ........ILAGPILILLILSMMVLPLPAFILDLLFTFNIALSIMVLLVAM
invA .....MLLSLLNSARLRPEL........LIL..VLMVMIISMFVIPLPTYLVDFLIALNIVLAILVFMGSF
ssaV .....MRSWLGEGVRAQQWLSVCAGRQDMVLATVLLIAIV.MMLLPLPTWMVDILITINLMFSVILLLIAI

          70        80        90       100       110       120       130
flhA FTQRTLDFAAFPTILLFTTLLRLALNVASTRIILMEGHTGAAAAGKVVEAFGHFLVGGNFAIGIVVFIILV
invA YIDRILSFSTFPAVLLITTLFRLALSISTSRLILIE.....ADAGEIIATFGQFVIGDSLAVGFVVFSIVT
ssaV YLSDPLDLSVFPSLLLITTLYRLSLTISTSRLVLLQ.....HNAGNIVDAFGKFVVGGNLTVGLVVFTIIT

        140       150       160       170       180       190       200
flhA IINFMVITKGAGRIAEVGARFVLDGMPGKQMAIDADLNAGLIGEDEAKKRRSEVTQEADFYGSMDGASKFV
invA VVQFIVITKGSERVAEVAARFSLDGMPGKQMSIDADLKAGIIDADAARERRSVLERESQLYGSFDGAMKFI
ssaV IVQFIVITKGIERVAEVSARFSLDGMPGKQMSIDGDLRAGVIDADHARTLRQHVQQESRFLGAMDGAMKFV

       210       220       230       240       250       260       270
flhA RGDAIAGILIMVINVVGGLLVGVLQHGMSIGSAAESYTLLTIGDGLVAQIPALVISTAAGVIVTRVSTD..
invA KGDAIAGIIIIFVNFIGGISVGMTRHGMDLSSALSTYTMLTIGDGLVAQIPALLIAISAGFIVTRVNGDS.
ssaV KGDTIAGIIVVLVNIIGGIIIAIVQYDMSMSEAVHTYSVLSIGDGLCGQIPSLLISLSAGIIVTRVPGEKR

        280       290       300       310       320       330
flhA QDVGEQMVGQLFSNPRVMLLAAAVLGLLGMVPGMPNLVFLLFTAALLGLAWWLRGRE.EKAPEEP......
invA DNMGRNIMTQLLNNPFVLVVTAILTISMGTLPGFPLPVFVILS.VVLSVLFYFKFREAKRSAAKPKTSKGE
ssaV QNLATELSSQIARQPQSLILTAVVLMLLALIPGFPFITLAFFS.ALLALPIILIRR..KKSVVSA......

     340      350       360       370       380       390       400
flhA QPVKMPENNSVVEATWNDVQLEDSLGMEVGYRLIPMVDFQQDGELLGRIRSIRKKFAQDMGFLPPVVHIRD
invA QPLSIEEKEGSSLGLIGDLDKVSTETVPLIL.LVPKSR.REDLEKAQLAERLRSQFFIDYGVRLPEVLLRD
ssaV NGVEAPEKDSMVPG...........ACPLILRLSPTLH...SADLIRDIDAMRWFLFEDTGVPLPEVNIEV

  410        420       430       440       450       460       470       480
flhA NMDLQPARYRILMKGVEIGSGDAYPGRWLAINPGTAAGTLPGEKTVDPAFGLDAIWIESALKEQAQIQGFT
invA GEGLDDNSIVLLINEIRVEQFTVYFDLMRVVNYSDEVVSFGINPTIHQQGSSQYFWVTHEEGEKLRELGYV
ssaV LPEPTEKLTVLLYQEPVFSLSIPAQADYLLIGADASVV...GDSQTLPNGMGQICWLTKDMAHKAQGFGLD

            490       500       510       520       530       540       550
flhA VVEASTVVATHLNHLIGQFSAELFGRQEAQQLLDRVSQEMPKLTEDLVPGVVTLTTLHKVLQNLLAEKVPI
invA LRNALDELYHCLAVTLARNVNEYFGIQETKHMLDQLEAKFPDLLKEVLRH.ATVQRISEVLQRLLSERVSV
ssaV VFAGSQRISALLKCVLLRHMGEFIGVQETRYLMNAMEKNYSELVKELQRQ.LPINKIAETLQRLVSERVSI

           560       570       580       590       600       610
flhA RDMRTILETLAEHAPLQSDPHELTAVVRVALGRAITQQWFPGNEEVQVIGLDTALERLLLQALQGGGG...
invA RNMKLIMEALALWAPREKDVINLVEHIRGAMARYICHKFANGGE.LRAVMVSAEVEDVIRKGIRQTSGSTF
ssaV RDLRLIFGTLIDWAPREKDVLMLTEYVRIALRRHILRRLNPEGKPLPILRIGEGIENLVRESIRQTAMGTY

     620       630       640       650       660       670       680
flhA ..LEPGLADRLLAQTQEALSRQEMLGAPPVLLVNHALRPLLSRFLRRSLPQLVVLSNLELSDNRHIRMTAT
invA LSLDPEASANLMDLITLKLDDLLIAHKDLVLLTSVDVRRFIKKMIEGRFPDLEVLSFGEIADSKSVNVIKT
ssaV TALSSRHKTQILQLIEQALKQ....SAKLFIVTSVDTRRFLRKITEATLFDVPILSWQELGEESLIQVVES

     690
flhA IGGK.........
invA I............
ssaV IDLSEEELADNEE
```
