## Extended Data Figure 6. Comparison of primary amino acid sequence of the SctN components of flagellar T3SS, SPI-1 and SPI-2 injectisomes. for "Flagellar system translocates T3SS effectors critical for *Salmonella* infection"

```
        1        10                  20        30        40
invC    MKTPRLLQYLA........YP...........QKITGPIIEAELRDVAIGELCEIRRGWH
fliI    .MTTRLTRWLTALDNFEAKMALLPAVRRYGRLTRATGLVLEATGLQLPLGATCIIERQDG
ssaN    .MKNELMQRLR......LKYPPPDGYCRWGRIQDVSATLLNAWLPGVFMGELCCIKPG..

                 50        60        70             80        90
invC    QKQVVARAQVVGLQRERTVLSLIGNAQGLSRDVVLYP.......TGRALSAWVGYSVLGA
fliI    PETKEVESEVVGFNGQRLFLMPLEEVEGILPGARVYARNGHGDGLQSGKQLPLGPALLGR
ssaN    ....EELAEVVGINGSKALLSPFTSTIGLHCGQQVMA.......LRRRHQVPVGEALLGR

             100       110       120       130       140       150
invC    VLDPTGKIVERFTPEVAPISEERVIDVAPP.SYASRVGVREPLITGVRAIDGLLTCGVGQ
fliI    VLDGGGKPLDGLP...APDTLETGALITPPFNPLQRTPIEHVLDTGVRAINALLTVGRGQ
ssaN    VIDGFGRPLDGRE...LPDVCWKDYDAMPP.PAMVRQPITQPLMTGIRAIDSVATCGEGQ

               160       170       180       190       200       210
invC    RMGIFASAGCGKTMLMHMLIEQTEADVFVIGLIGERGREVTEFVDMLRASHKKEKCVLVF
fliI    RMGLFAGSGVGKSVLLGMMARYTRADVIVVGLIGERGREVKDFIENILGPDGRARSVVIA
ssaN    RVGIFSAPGVGKSTLLAMLCNAPDADSNVLVLIGERGREVREFIDFTLSEETRKRCVIVV

               220       230       240       250       260       270
invC    ATSDFPSVDRCNAAQLATTVAEYFRDQGKRVVLFIDSMTRYARALRDVALASGERPARRG
fliI    APADVSPLLRMQGAAYATRIAEDFRDRGQHVLLIMDSLTRYAMAQREIALAIGEPPATKG
ssaN    ATSDRPALERVRALFVATTIAEFFRDNGKRVVLLADSLTRYARAAREIALAAGETAVSGE

               280       290        300       310       320       330
invC    YPASVFDNLPRLLERPGAT..SEGSITAFYTVLLESEEEADPMADEIRSILDGHLYLSRK
fliI    YPPSVFAKLPALVERAGNGIHGGGSITAFYTVLTEGDDQQDPIADSARAILDGHIVLSRR
ssaN    YPPGVFSALPRLLERTGMG..EKGSITAFYTVLVEGDDMNEPLADEVRSLLDGHIVLSRR

                   340       350       360       370       380       390
invC    LAGQGHYPAIDVLKSVSRVFGQVTTPTHAEQASAVRKLMTRLEELQLFIDLGEYRPGENI
fliI    LAEAGHYPAIDIEASISRAMTALITEQHYARVRLFKQLLSSFQRNRDLVSVGAYAKGSDP
ssaN    LAERGHYPAIDVLATLSRVFPVVTSHEHRQLAAILRRCLALYQEVELLIRIGEYQRGVDT

                   400       410       420       430
invC    DNDRAMQMRDSLKAWLCQPVAQYSSFDDTLSGMNAFADQN
fliI    MLDKAITLWPQLEAFLQQGIFERADWEDSLQALDLIFPTV
ssaN    DTDKAIDTYPDICTFLRQSKDEVCGPELLIEKLHQILTE.
```
