## Appendix A. Code for detection of effector translocation start from raw luminescent data. for "Flagellar system translocates T3SS effectors critical for *Salmonella* infection"

import pandas as pd

import numpy as np

import matplotlib.pyplot as plt

from scipy.optimize import curve_fit

file_name = 'data.xlsx'

df = pd.read_excel(file_name)

time = df.iloc[:, 0].values

protein_cols = df.columns[1:]

def gompertz_rise(t, lower, upper, k, ti):

"""Asymmetric growth model."""

return lower + (upper - lower) * np.exp(-np.exp(-k * (t - ti)))

def analyze_protein_robust(x, y):

\# Step 1: Find Peak

peak_idx = np.argmax(y)

if peak_idx < 3: # If peak is at the very beginning

return None

x_rise = x[:peak_idx + 2]

y_rise = y[:peak_idx + 2]

\# Step 2: Internal Normalization to 0-1 (Crucial for fit stability)

y_min, y_max = np.min(y_rise), np.max(y_rise)

if y_max == y_min: return None

y_norm = (y_rise - y_min) / (y_max - y_min)

\# Step 3: Set Initial Guesses and Strict Bounds

\# [lower, upper, k, ti]

p0 = [0, 1, 1.0, np.median(x_rise)]

bounds = ([-0.5, 0.8, 0.001, x_rise[0]], [0.5, 1.5, 100, x_rise[-1]])

try:

\# Step 4: Fit the model

popt, _ = curve_fit(gompertz_rise, x_rise, y_norm, p0=p0, bounds=bounds, maxfev=20000)

lower_n, upper_n, k, ti = popt

\# Step 5: Calculate 'Ct' Onset in Normalized Space

onset_time = ti - (np.log((3 + np.sqrt(5)) / 2) / k)

\# Convert back to original scale for plotting

y_fit_norm = gompertz_rise(x_rise, *popt)

y_fit_orig = y_fit_norm * (y_max - y_min) + y_min

return {

'Onset_Time': max(x[0], onset_time), # Clip to start of experiment

'Peak_Time': x[peak_idx],

'Peak_Intensity': y_max,

'Growth_Rate': k,

'x_rise': x_rise,

'y_fit': y_fit_orig

}

except Exception as e:

return None

\# --- Main Analysis Loop ---

all_results = []

for col in protein_cols:

signal = df[col].values

res = analyze_protein_robust(time, signal)

if res:

all_results.append({

'Protein_ID': col,

'Onset_Time_Ct': round(res['Onset_Time'], 3),

'Peak_Time': round(res['Peak_Time'], 3),

'Peak_Intensity': round(res['Peak_Intensity'], 4),

'Growth_Rate': round(res['Growth_Rate'], 4)

})

\# Plotting

plt.figure(figsize=(8, 4))

plt.plot(time, signal, 'ko', markersize=2, alpha=0.3, label='Raw Data')

plt.plot(res['x_rise'], res['y_fit'], 'r-', linewidth=2, label='Rise Fit')

plt.axvline(x=res['Onset_Time'], color='blue', linestyle='--', label=f"Onset: {res['Onset_Time']:.2f}")

plt.title(f"Analysis: {col}")

plt.legend()

plt.show()

else:

\# Fallback for failed fits

print(f"Fit failed for {col}. Recording N/A.")

all_results.append({

'Protein_ID': col, 'Onset_Time_Ct': 'FIT_FAILED',

'Peak_Time': 'N/A', 'Peak_Intensity': 'N/A', 'Growth_Rate': 'N/A'

})

results_df = pd.DataFrame(all_results)

output_file = "Translocation_Analysis.xlsx"

results_df.to_excel(output_file, index=False)

print(f"\nDone! Results saved to {output_file}")

print(results_df)
